# Land-use history impacts spatial patterns and composition of woody plant species across a 35-hectare temperate forest plot

**DOI:** 10.1101/2021.04.07.438791

**Authors:** D.A. Orwig, J.A. Aylward, H.L. Buckley, B.S. Case, A.M. Ellison

**Affiliations:** Harvard Forest, Harvard University, Petersham, MA 01366 USA; School of Science, Auckland University of Technology, Private Bag 92006, Auckland, 1142, New Zealand

**Keywords:** ForestGEO, Harvard Forest, land-use history, spatial point-pattern analysis, temperate forest, *Tsuga canadensis*

## Abstract

Land-use history is the template upon which contemporary plant and tree populations establish and interact with one another and exerts a legacy on the structure and dynamics of species assemblages and ecosystems. We use the first census (2010–2014) of a 35-ha forest-dynamics plot at the Harvard Forest in central Massachusetts to explore such legacies. The plot includes 108,632 live stems ≥ 1 cm in diameter (2215 individuals/ha) and 7,595 dead stems ≥ 5 cm in diameter. Fifty-one woody plant species were recorded in the plot, but two tree species— *Tsuga canadensis* (eastern hemlock) and *Acer rubrum* (red maple)—and one shrub— *Ilex verticillata* (winterberry)—comprised 56% of all stems. Live tree basal area averaged 42.25 m^2^/ha, of which 84% was represented by *T. canadensis* (14.0 m^2^/ ha), *Quercus rubra* (northern red oak; 9.6 m^2^/ ha), *A. rubrum* (7.2 m^2^/ ha) and *Pinus strobus* (eastern white pine; 4.4 m^2^/ ha). These same four species also comprised 78% of the live aboveground biomass, which averaged 245.2 Mg/ ha, and were significantly clumped at distances up to 50 m within the plot. Spatial distributions of *A. rubrum* and *Q. rubra* showed negative intraspecific correlations in diameters up to at least a 150-m spatial lag, likely indicative of competition for light in dense forest patches. Bivariate marked point-pattern analysis showed that *T. canadensis* and *Q. rubra* diameters were negatively associated with one another, indicating resource competition for light. Distribution and abundance of the common overstory species are predicted best by soil type, tree neighborhood effects, and two aspects of land-use history: when fields were abandoned in the late 19^th^ century and the succeeding forest types recorded in 1908. In contrast, a history of intensive logging prior to 1950 and a damaging hurricane in 1938 appear to have had little effect on the distribution and abundance of present-day tree species.

## Introduction

In forested landscapes around the world, legacies of human activities have shaped the composition, size structure, and spatial patterns of trees, understory vegetation, and associated ecosystem processes (Birks et al. 1988, Turner et al. 1990, Russell 1997, Foster and Aber 2004, Ellison et al. 2014). The extent of the interactions between anthropogenic effects and abiotic factors such as climate, soils, and episodic disturbances in shaping vegetation patterns depends on the intensity of the effects and the spatial scale of analysis (Rackham 1986, Glitzenstein et al. 1990, Zimmerman et al. 1995). A complex interplay of succession, competition, disturbance, environment, and land use shape dynamics and patterns of forests at local-to-regional scales (Condit et al. 2000, Thompson et al. 2002, Chazdon 2003, Van Gemerden et al. 2003).

The forests of New England, USA have been shaped by a variety of natural and anthropogenic factors. As in other forests, the geology and climate of New England define the broad patterns of current forest composition (Foster et al. 1992, Hall et al. 2002), but the shifts in species abundance and distribution patterns that have occurred since Europeans colonized New England more than 400 years ago have resulted in a relatively homogenous assemblage of young, even-aged stands with fewer late-successional species (Thompson et al. 2013). In Massachusetts, modern vegetation exhibits only weak relationships to broad climatic gradients because of the overwhelming influence of past land use (Foster et al. 1998). An increasing emphasis in ecological studies is evaluating the relative importance of historic land-clearing, agriculture, intensive harvesting (Foster 1992, Thompson et al. 2002, Hogan et al. 2016), and natural episodic storms (Foster and Boose 1992, Zimmerman et al. 1995) on current-day structure and species composition of forest stands (Motzkin et al. 1996, Motzkin et al. 1999).

Harvard Forest is an ideal location to investigate how spatial patterns and composition of woody plant are influenced by land-use history impacts. For more than a century, HF researchers have investigated impacts of land-use on forests and how New England’s forests are continuing to change as the regional climate changes, populations of large herbivores wax and wane, and nonnative insects and pathogens establish, irrupt, and kill tree species (Foster and Aber 2004).

Here, we describe the results of the first census of a 35-ha forest-dynamics plot at the Harvard Forest and examine how its structure and composition relates to interactions between land-use history and ecological processes. We first describe the composition and structure of the woody plants in this plot and assess spatial associations within and among the dominant species using univariate and bivariate spatial point-pattern analysis. Second, we uncover the influence of historical land-use and natural disturbances on the current-day structure and composition of this forest plot. We pay particular attention to patterns of distribution and abundance of *Tsuga canadensis* (eastern hemlock) and its relationship to other species in the plot because previous work has shown it to be a foundation species in this forest (sensu (Ellison 2019)). *Tsuga canadensis* is currently threatened and declining throughout much of its range due to a nonnative insect, *Adelges tsugae* (hemlock woolly adelgid; HWA) and its decline and loss are likely to have profound impacts on forest structure and composition (Orwig et al. 2013, Foster 2014).

## Methods

### Site description

The 35-ha (500 × 700 m) forest-dynamics plot at Harvard Forest (HF), is part of a global network of Forest Global Earth Observatory (ForestGEO) plots established to monitor, understand, and predict forest dynamics and responses to global change (Anderson-Teixeira et al. 2015). The HF ForestGEO plot (southwest corner at 42.5386 °N, 72.1798 °W) is located within the 380-ha HF Prospect Hill tract in Petersham, Massachusetts, USA within the Worcester/Monadnock Plateau ecoregion (Griffith et al. 1994) of Transition Hardwoods-White Pine-Hemlock forests (Westveld 1956) (Fig. 1). Elevations in the plot range from 340.2 to 367.8 m a.s.l. Soils include Gloucester stony loam, Acton stony loam and Whitman very stony silt loams, all of which are gravelly and fine sandy loam soils that developed in glacial tills overlying gneiss and schist bedrock (Simmons 1941). The north-central portion of the plot contains a 3-ha peat swamp with muck soils that has been colonized at intervals by *Castor canadensis* (beaver). Average (1964-2019) annual temperature at the site is 7.9 °C and the annual precipitation of 1090 mm is distributed evenly throughout the year (Boose and Gould 2019).

**Figure 1.**
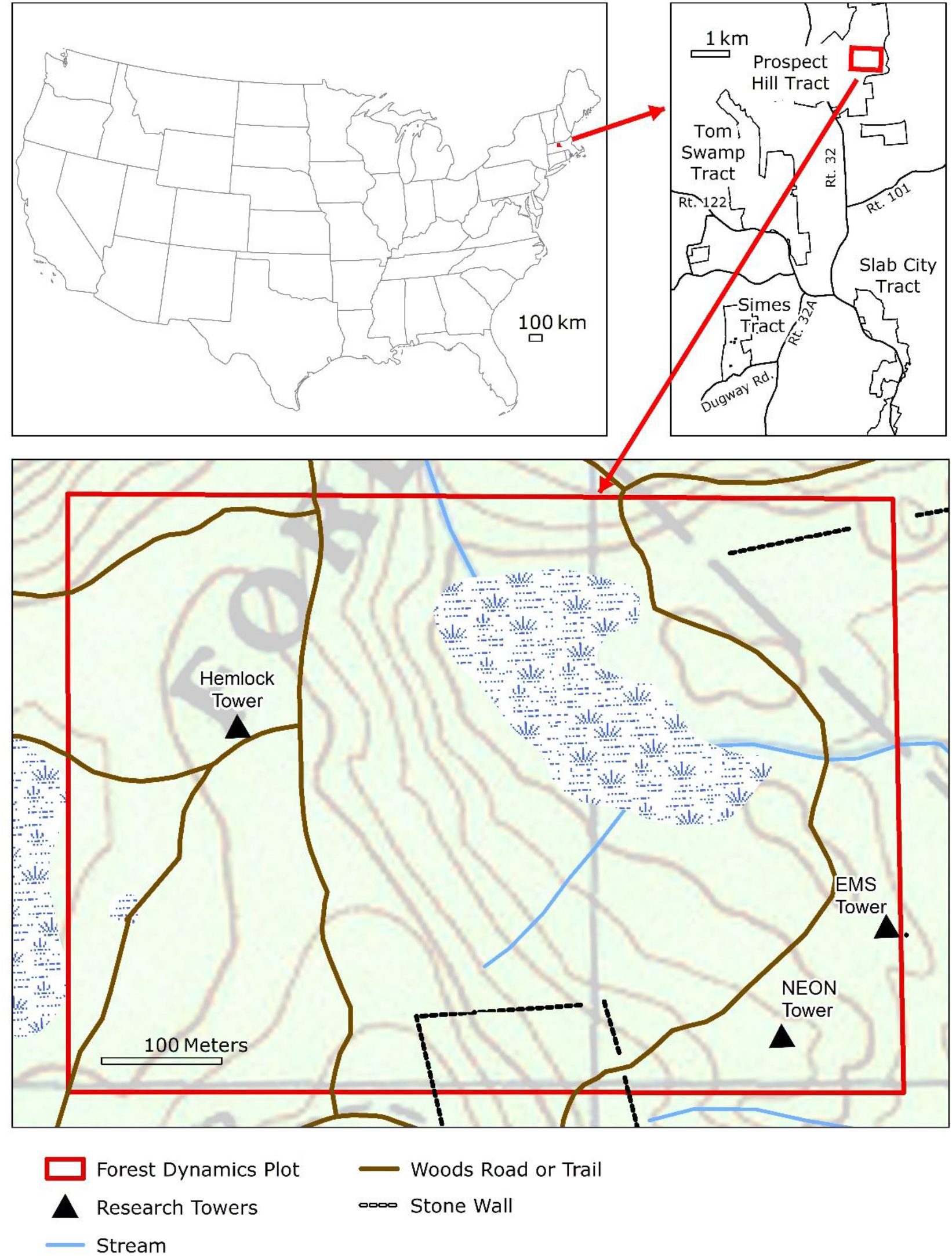
The 500 x 700 m ForestGEO plot located in the town of Petersham, MA on the Prospect Hill tract of HF (upper right panel), showing locations of three eddy-flux towers (that measure net ecosystem exchange of carbon and water between the atmosphere and the ecosystem), old forest roads, stone walls (denoted by dotted lines), and the central swamp area, superimposed on topographic contour lines (lower panel).

### Land-use history

We examined the influence of past land-use history (derived from forest stand descriptions of dates of field abandonment, areas used as woodlot, pasture, or cultivation; presence of distinct plow horizon; silviculture treatments; and salvage operations), historical events (e.g., insect outbreaks, storms and associated degree of forest damage (Rowlands 1941)), and biophysical attributes (roads, soil type, slope, aspect, elevation, and distance to streams) on current forest composition and species distribution within the plot by using data from the document archives at HF (http://harvardforest.fas.harvard.edu/document-archive). Original maps of activity were manually transcribed to standardized base maps and then scanned and digitized as shapefiles in ArcView GIS 3.2. The shapefiles were then transformed to Massachusetts State Plane Meters (NAD83 projection) in ArcGIS to align better with aerial photographs and linear features (trails, stonewalls, etc.) downloaded from MassGIS (Hall 2005) and used in spatial analyses (see below).

Pollen evidence suggests that prior to European settlement, Prospect Hill was a mixture of old-growth northern hardwoods, *T. canadensis*, and *Pinus strobus* (eastern white pine) (Foster 1992, Foster et al. 1992). Following European arrival, the site then experienced complex ownership and intensive land-use over the next few centuries, both of which are largely representative of the New England region (Ellison et al. 2014). Forest clearing began in 1750 and reached a maximum in the 1840s, by which time close to 80% of the original forests had been cleared for agriculture (Fisher 1933, Raup and Carlson 1941). Field abandonment began in 1850 and continued through 1905 in the southern half of the plot (Fig. 2a). Reforestation of those fields continued through the 20th century (Foster 1992). The western, northern, and northeastern areas of the plot remained permanently wooded, but experienced various types of selective cutting in the 1790s and 1870s (Foster 1992). The first maps characterizing forest types of individual stands were completed in 1908 and classified the permanent woodlots in the western third of the plot as being comprised of hardwoods, white pine-hardwoods, hemlock, and red maple (Fig 2b). Many *Castanea dentata* (American chestnut) died in 1912–1914 from infection by *Endothia parasitica* (chestnut blight) (McLachlan et al. 2000) and forests were damaged by natural disturbances including an ice storm in 1921 and one of the most damaging hurricanes to hit New England in 1938. The hurricane and subsequent salvage logging resulted in the loss of as much as 70% of the standing timber on HF properties (Foster and Boose 1992).

**Figure 2.**
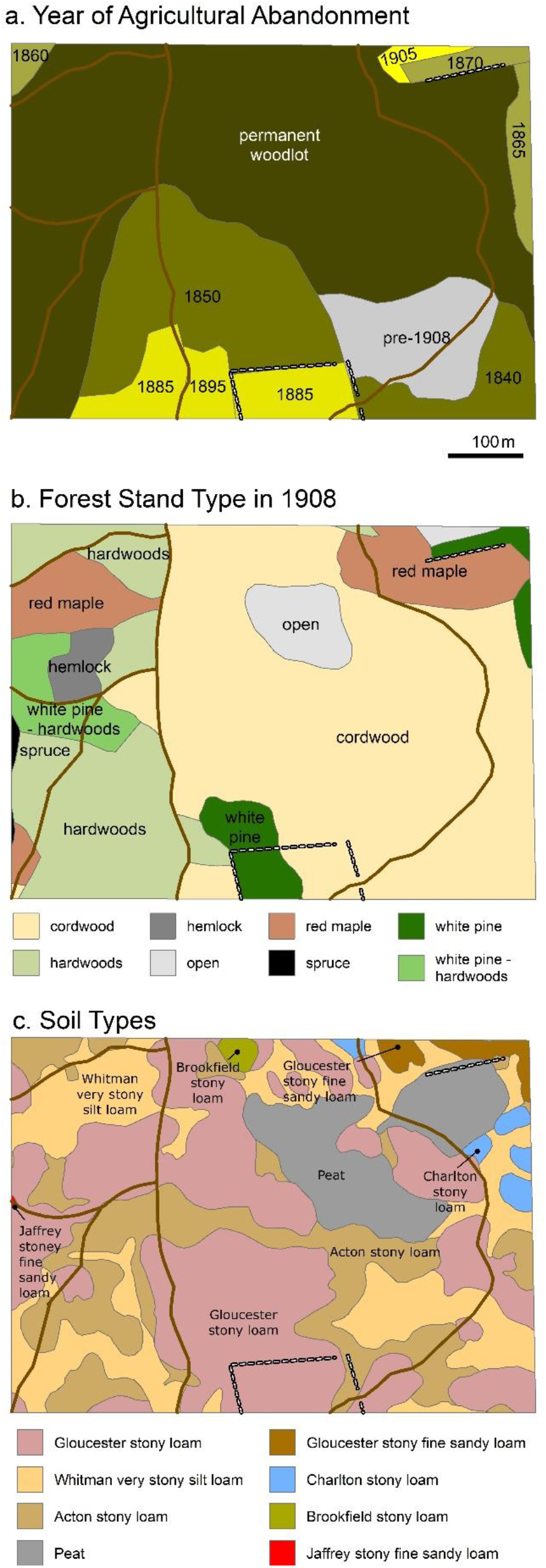
Location of a) historical fields and their agricultural date of abandonment, b) forest stands as described in 1908, and c) soil type within the HF ForestGEO plot. GIS layers obtained from Harvard Forest Document Archive HF 110.

The central sections of the plot, containing mostly stony loam soils and no visible signs of a plow layer, were unimproved pastures abandoned in the mid-19th century (Motzkin et al. 1999)(Fig 2c). These areas reforested and were classified as cordwood (poor hardwood) in 1908 (Fig 2b), except for an area classified as open, which is the beaver swamp. Much of the cordwood section was subsequently clear-cut in the 1920s and then thinned or salvaged in the late 1940s following the 1938 hurricane. *Pinus resinosa* (red pine) and *Picea abies* (Norway spruce) plantations were established in portions of these abandoned pastures in the mid-1920s and early 1930s. The southcentral area of the plot contained areas of improved pasture and cultivation (Motzkin et al. 1999) and was classified as containing white pine in 1908. This area was clear-cut in the 1920s and a portion of it was clear-cut again in 1980, resulting in many small diameter, multi-stemmed trees. Additional biotic changes that impacted the plot included the exotic *Lymantria dispar* (gypsy moth), which lead to widespread defoliation of hardwoods during 1944–45 and 1981; *Cryptococcus fagisuga* (beech scale insect) combined with *Neonectria* fungal spp. (beech-bark disease), which has led to the decline and death of larger *Fagus grandifolia* (American beech); and *Adelges tsugae*, which was first observed in the plot in 2008, rapidly spread throughout the plot, subsequently killing hundreds of *T. canadensis* stems and threatening the rest (Orwig et al. 2018).

### Plot establishment and woody stem census

During March 2010, professional surveyors delineated the plot boundaries, established a continuous grid of 20 × 20-m quadrats, and measured the elevation at each post using a Sokkia SET600 Total Station (Olathe, Kansas, USA). During the summers of 2010 and 2011, all woody stems ≥ 1 cm in diameter at breast height (DBH; 1.3 m above the ground level) were uniquely tagged, identified (nomenclature follows (Haines 2011)), and measured to the nearest 0.1 cm (Condit 1998). All dead stems ≥ 5 cm diameter that were standing and > 45 degrees from horizontal also were tagged, identified, and measured. The swamp located in the center of the plot was sampled when the ground was frozen during the winter months of 2012–2014. Each tagged stem was mapped within one of four 10 × 10 m subquadrats within each quadrat on a scale-drawn map data sheet. Each map was then scanned and individual stems were digitized using the Image J processing program (Rasband 2012), and converted to local (*x, y*) coordinates within a quadrat using R (v.3.6.1) (R Core Team 2013) and the CTFS R package (Condit 2014).

### Forest species composition and stand structure

Estimates of stem densities were derived from total counts in which multi-stemmed individuals were considered as a single stem, whereas estimates of basal area and biomass were derived from the sum of all stems ≥ 1 cm DBH (Gilbert et al. 2010). Biomass of living woody stems was estimated from DBH using allometric equations (Table S1).

### Spatial analysis

We assessed the spatial patterns of the seven most abundant tree species across the entire plot using the pair-correlation function (*g*(*r*);(Wiegand and Moloney 2014)), for which the value of the function represents the degree of clustering (*g*(*r*) > 1) or overdispersion (*g*(*r*) < 1) at a given spatial lag (distance between neighboring trees). We compared the observed pair-correlation statistic to that expected if trees were distributed randomly (*g*(*r*) = 1) within the plot using 199 Monte Carlo CSR (complete spatial randomness) simulations of the tree map for each species.

To test for the effects of intraspecific competition we used the univariate mark-correlation function (*kmm*(*r*); (Wiegand and Moloney 2004, Wiegand and Moloney 2014)) to test whether the size of each of the seven most abundant tree species depended on its proximity to neighbors of its own species. The value of *kmm*(*r*) represents the relative sizes of trees at a given spatial lag and indicates if trees are larger or smaller than expected at a given spatial lag. We compared the observed univariate mark-correlation function statistic to that expected if the sizes of trees were randomly assigned across individuals using 199 Monte Carlo simulations for each species, i.e., the spatial pattern of the trees remained the same, but their sizes were shuffled (Jacquemyn et al. 2010). Spatial analyses were not conducted on shrub species as many only occurred in the central swamp area.

Prior work has shown that the shade-tolerant *T. canadensis* is an important foundation tree species, creating and strongly controlling the microenvironment, understory vegetation, and ecosystem dynamics (Ellison et al. 2005, Orwig et al. 2013). Thus, we assessed the potential influence of *T. canadensis* on the sizes of each of the other most common tree species in the plot using a bivariate marked point pattern analysis (Schlather’s version of Moran’s I mark-correlation function (*Im1m2*(*r*); (Wiegand and Moloney 2014)). This statistic determines if tree sizes are spatially correlated: individuals are smaller or larger than expected at various distances from a neighbor. We compared the observed *Im1m2*(*r*) to that expected if the sizes of trees were randomly assigned across individuals using 199 Monte Carlo simulations for each species (Jacquemyn et al. 2010). All spatial pattern analyses were performed using the 2018 version of the software Programita (Wiegand and Moloney 2004, Wiegand and Moloney 2014).

GIS overlays of past land use, historical events, and biophysical attributes were used as covariates in a conditional inference regression-tree model to predict diameter and abundance of the most common overstory species in the plot (Table 1). Using the ‘cforest’ function in the R package ‘party’ (Version 1.3-5) (Hothorn et al. 2013) the outcomes of 500 conditional inference tree models (Hothorn et al. 2006) were compiled and the relative importance of explanatory variables were ranked across all models. The conditional inference algorithm is based on a random forest machine-learning algorithm (Breiman 2001) used in many ecological modeling contexts (e.g., (Fox et al. 2017, Mi et al. 2017, Mohapatra et al. 2019, Shearman et al. 2019)).

**Table 1.** Description of land-use history, disturbance, stand, and biophysical variables converted to GIS shapefiles and used to predict current tree species abundance and DBH values across the Harvard Forest ForestGEO plot.

| Predictor | Description |
| --- | --- |
| *Land use* |  |
| Stand type – 1908, 1947, 1986 | Early forest stand descriptions in plot recorded by forest type and year |
| Allen Land Use | Land-use descriptions derived from degree of soil disturbance, including plow (Ap) horizon presence and depth, recorded by previous HF soil scientist, Arthur Allen. |
| Field abandonment | Years since the date of field abandonment |
| 20<sup>th</sup> C. Salvage cutting | Areas that experienced cutting following wind damage or other natural disturbance in the early to mid-1900s |
| 20<sup>th</sup> C. intensive cutting | Areas that experienced clearcut, shelterwood or reproduction cuts during the early to mid-1900s |
| *Natural disturbance* |  |
| Hurricane damage | Data collected between 1939-1941 on degree of overstory trees uprooted, leaning or broken after 1938 hurricane (Rowlands 1941). |
| *Stand features* |  |
| Mean DBH of trees within 10m | Mean DBH of trees within 10m of individual tree stem |
| CV DBH of trees within 10m | Coefficient of variation of DBH of trees within 10m |
| Number of trees within 10m | Number of trees within 10m of individual tree stem |
| Mean distance to trees within 10m | Mean distance to trees within 10m of individual tree stem |
| CV distance to trees within 10m | Coefficient of variation in distance to trees within 10m of individual tree stem |
| *Biophysical features* |  |
| Elevation | Elevation of quadrat as determined from NASA Goddard's Lidar, hyperspectral and thermal (G-LiHT) airborne imager. |
| Distance to streams (m) | Distance from individual tree stem to streams as identified by the National Hydrography Dataset |
| Soil drainage class | USDA Natural Resources Conservation Service Soil Survey Geographic (SSURGO) database soil attribute |
| Simmons soil type | Soil Classification from 1:24000 scale surveys Simmons (1941) |

The conditional inference method improves on the variable ranking methodology by applying a permutation importance algorithm that corrects for variable selection bias resulting from a mix of categorical and continuous explanatory variables that are correlated to varying degrees or that have complex interactions (Strobl et al. 2007). Variable importance scores are calculated by determining the marginal loss of prediction accuracy from any given model iteration after removing each explanatory variable. Overall variable importance is determined by averaging the variable-wise decrease in accuracy scores over all 500 model iterations and using this to rank the overall importance of each variable across all models. Species-specific abundances or sizes were predicted for each of the seven most abundant overstory species conditional on their observed locations. A moving-window focal analysis of the count of trees for each species in a 20-m rectangle around each tree’s location generated relative abundance (stems/ha). Then, given the location of a tree, relative abundance was sampled from the species-specific raster using an interpolation function to compute the average relative abundance around that location.

### Data availability

Data associated with this study are publicly available from the HF data archive (Orwig et al. 2015): HF253. http://harvardforest.fas.harvard.edu.

## Results

### Composition and stand structure

Within the 35-ha HF ForestGEO plot, we identified 108,632 live stems ≥ 1cm DBH, representing 77,536 individuals (2215 ha^-1^) of 51 woody species in 17 families (Table S2).

Common families were Betulaceae, Rosaceae, and Pinaceae (six species each), and Fagaceae and Adoxaceae (five species each). Four tree species (*T. canadensis*, *Acer rubrum* [red maple], *Q. rubra*, and *P. strobus*) and one shrub, *Ilex verticillata* (winterberry), accounted for 63% of all stems (Table 2). Live tree basal area was 42.25 m^2^/ha and live aboveground biomass was 245.2 Mg/ha. Eighty-four percent of the basal area and 78% of the biomass was represented by *T. canadensis* (14.0 m^2^/ha; 61.1 Mg/ha), *Q. rubra* (9.6 m^2^/ha; 75.1 Mg/ha), *A. rubrum* (7.2 m^2^/ha; 33.8 Mg/ha) and *P. strobus* (4.4 m^2^/ha; 20.7 Mg/ha). The live tree diameter distributions of *T. canadensis* and *F. grandifolia* were strongly right-skewed (reverse-J shaped), whereas those of *A. rubrum*, *Q. rubra*, *P. strobus, Betula lenta* (black birch), and *B. alleghaniensis* (yellow birch) were less right-skewed (Fig. 3). Seventy-seven live stems of *Betula*, *Picea*, and *Quercus* could not be identified to species, mostly due to difficulties of differentiating between young *Betula* saplings and between *Quercus rubra* (northern red oak) and *Q. velutina* (black oak).

**Table 2.** List of total live woody plant density, basal area, and biomass within the 35 ha HF ForestGEO plot in 2014.

| Scientific name | Total live tree Density<br>(35 ha<sup>-1</sup>) | Total live Basal<br>area (m<sup>2</sup>) | Total live Biomass<br>(Mg ) |
| --- | --- | --- | --- |
| *Acer pensylvanicum* | 339 | 0.59 | 1.13 |
| *Acer rubrum* | 9,723 | 253.54 | 1182.86 |
| *Acer saccharum* | 1 | 3.12e-03 | 0.02 |
| *Alnus incana* | 479 | 0.68 | 0.60 |
| *Amelanchier laevis* | 572 | 0.35 | 0.61 |
| *Aronia melanocarpa* | 413 | 0.07 | 0.10 |
| *Betula alleghaniensis* | 4,059 | 36.96 | 207.73 |
| *Betula lenta* | 1,430 | 21.14 | 124.04 |
| *Betula papyrifera* | 537 | 14.80 | 72.76 |
| *Betula populifolia* | 108 | 1.49 | 7.18 |
| *Castanea dentata* | 732 | 1.12 | 4.35 |
| *Crataegus spp.* | 180 | 0.14 | 0.27 |
| *Fagus grandifolia* | 3,802 | 20.93 | 138.58 |
| *Frangula alnus* | 3 | 7.42e-04 | 4.90e-04 |
| *Fraxinus americana* | 186 | 3.84 | 23.73 |
| *Fraxinus nigra* | 34 | 0.17 | 0.82 |
| *Hamamelis virginiana* | 1,931 | 3.10 | 5.77 |
| *Ilex laevigata* | 2 | 1.39e-03 | 2.76e-03 |
| *Ilex mucronata* | 598 | 0.64 | 0.58 |
| *Ilex verticillata* | 9,874 | 3.62 | 6.15 |
| *Juniperus communis* | 1 | 4.52e-04 | 4.20e-04 |
| *Kalmia latifolia* | 3,914 | 3.27 | 7.64 |
| *Lindera benzoin* | 66 | 0.02 | 0.04 |
| *Lyonia ligustrina* | 1,178 | 0.41 | 2.04 |
| *Nyssa sylvatica* | 180 | 2.63 | 11.25 |
| *Ostrya virginiana* | 24 | 0.06 | 0.19 |
| *Picea abies* | 900 | 24.43 | 93.11 |
| *Picea rubens* | 101 | 3.61 | 15.15 |
| *Pinus resinosa* | 790 | 67.23 | 330.28 |
| *Pinus strobus* | 2,126 | 155.68 | 724.64 |
| *Populus grandidentata* | 2 | 0.03 | 0.14 |
| *Populus tremuloides* | 1 | 0.01 | 0.05 |
| *Prunus pensylvanica* | 11 | 0.05 | 0.98 |
| *Prunus serotina* | 250 | 5.48 | 34.85 |
| *Quercus alba* | 38 | 1.89 | 14.53 |
| *Quercus rubra* | 3,896 | 334.99 | 2,627.07 |
| *Quercus velutina* | 206 | 19.28 | 164.46 |
| *Rhododendron prinophyllum* | 127 | 0.05 | 0.25 |
| *Salix spp.* | 2 | 1.59e-04 | 1.50e-03 |
| *Sambucus racemosa* | 2 | 5.65e-04 | 4.03e-03 |
| *Sorbus Americana* | 66 | 0.26 | 2.78 |
| *Toxicodendron radicans* | 1 | 1.13e-04 | 1.05e-04 |
| *Toxicodendron vernix* | 521 | 0.32 | 0.38 |
| *Tsuga canadensis* | 22,880 | 491.07 | 2138.00 |
| *Ulmus Americana* | 1 | 2.84e-04 | 3.85e-04 |
| *Vaccinium corymbosum* | 3,531 | 2.39 | 9.58 |
| *Viburnum acerifolium* | 39 | 0.01 | 0.07 |
| *Viburnum dentatum* | 325 | 0.08 | 0.52 |
| *Viburnum lantanoides* | 75 | 0.01 | 0.01 |
| *Viburnum nudum* | 1,182 | 0.44 | 2.27 |

**Figure 3.**
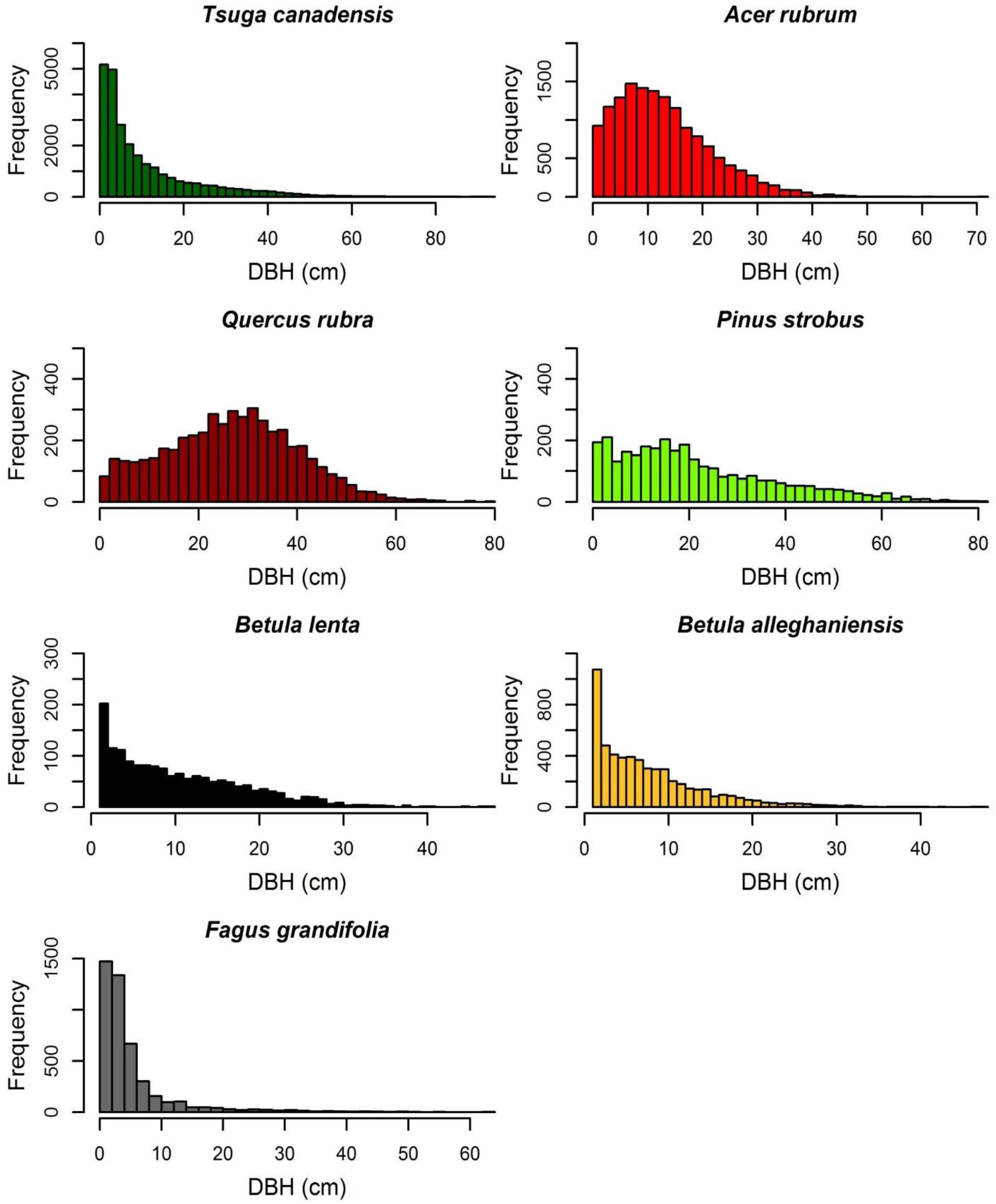
Diameter distribution of the seven most common overstory species within the HF ForestGEO plot.

In contrast, 73% of tagged stems and 69% of live individuals within the plot were < 10 cm DBH (Fig. 4). These same stems comprised only 5% of the total live plot basal area and 3% of the total live plot biomass (Table 2). Shrub species made up many of these stems and included *I. verticillata*, *Vaccinium corymbosum* (highbush blueberry), and *Kalmia latifolia* (mountain laurel). Nonnative species in the plot included 1687 stems of *Picea abies* (Norway spruce) and *Pinus resinosa* (red pine) that remained from early 20^th^-century conifer plantings and three stems of *Frangula alnus* (glossy false buckthorn). Ten species had only one or two stems within the plot (Table 2). Finally, there were 7595 dead stems ≥ 5 cm DBH within the plot, > 50% of which were *T. canadensis*, *P. strobus*, or *A. rubrum*. Dead tree basal area was 4.18 m^2^/ha and dead aboveground biomass was 17.53 Mg/ha.

**Figure 4.**
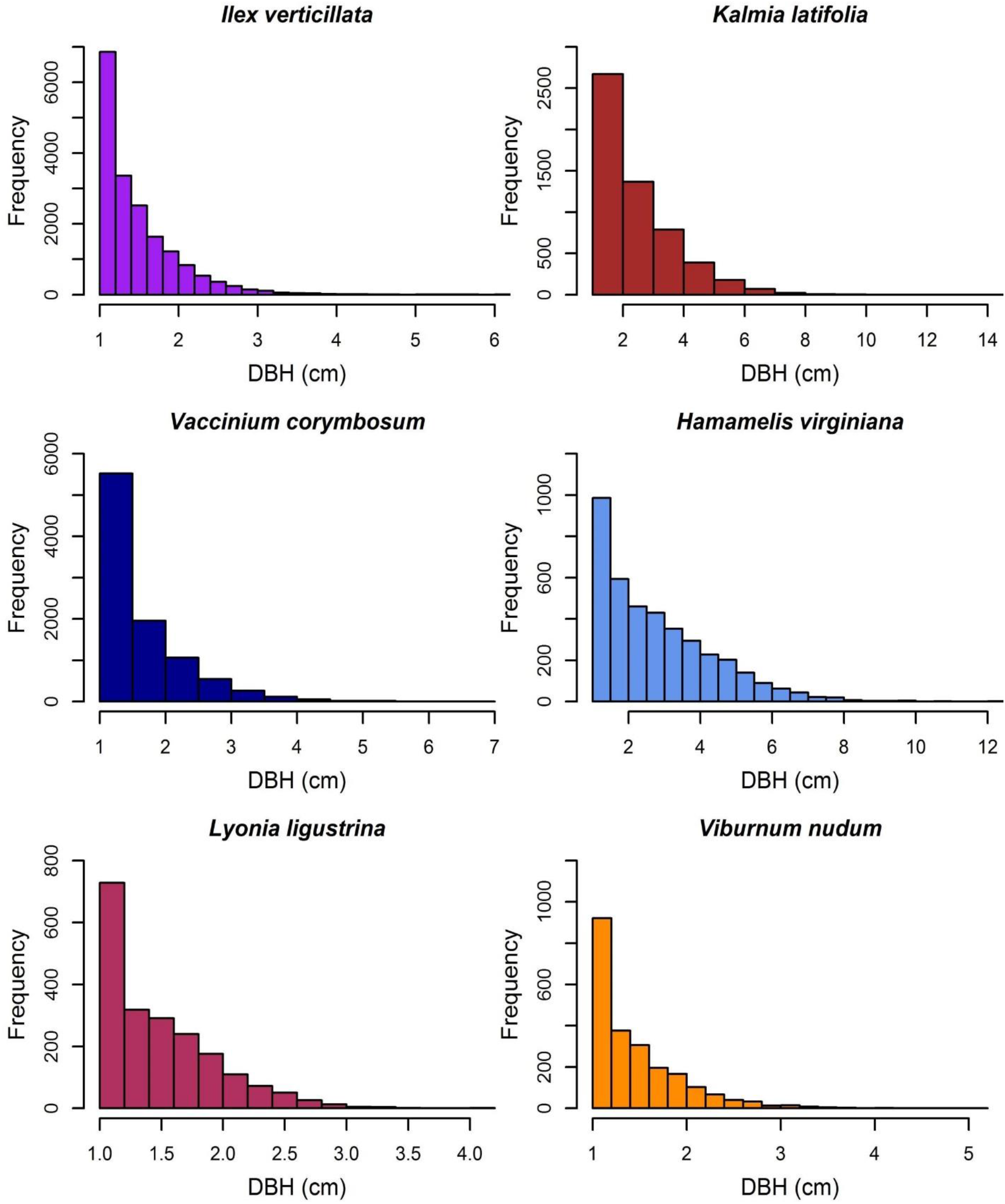
Diameter distribution of the six most common understory species within the HF ForestGEO plot.

### Spatial structure related to past land-use impacts

The spatial distributions of the seven most common species varied across the plot (Fig. 5). *Pinus strobus* was common throughout the plot. *Tsuga canadensis* was most abundant in the western, northern and eastern portions of the plot, whereas *Q. rubra* and *A. rubrum* dominated the central and southern areas. Both *Betula* species were most abundant in the central and eastern sections, and *F. grandifolia* was most common in the southeastern section.

**Figure 5.**
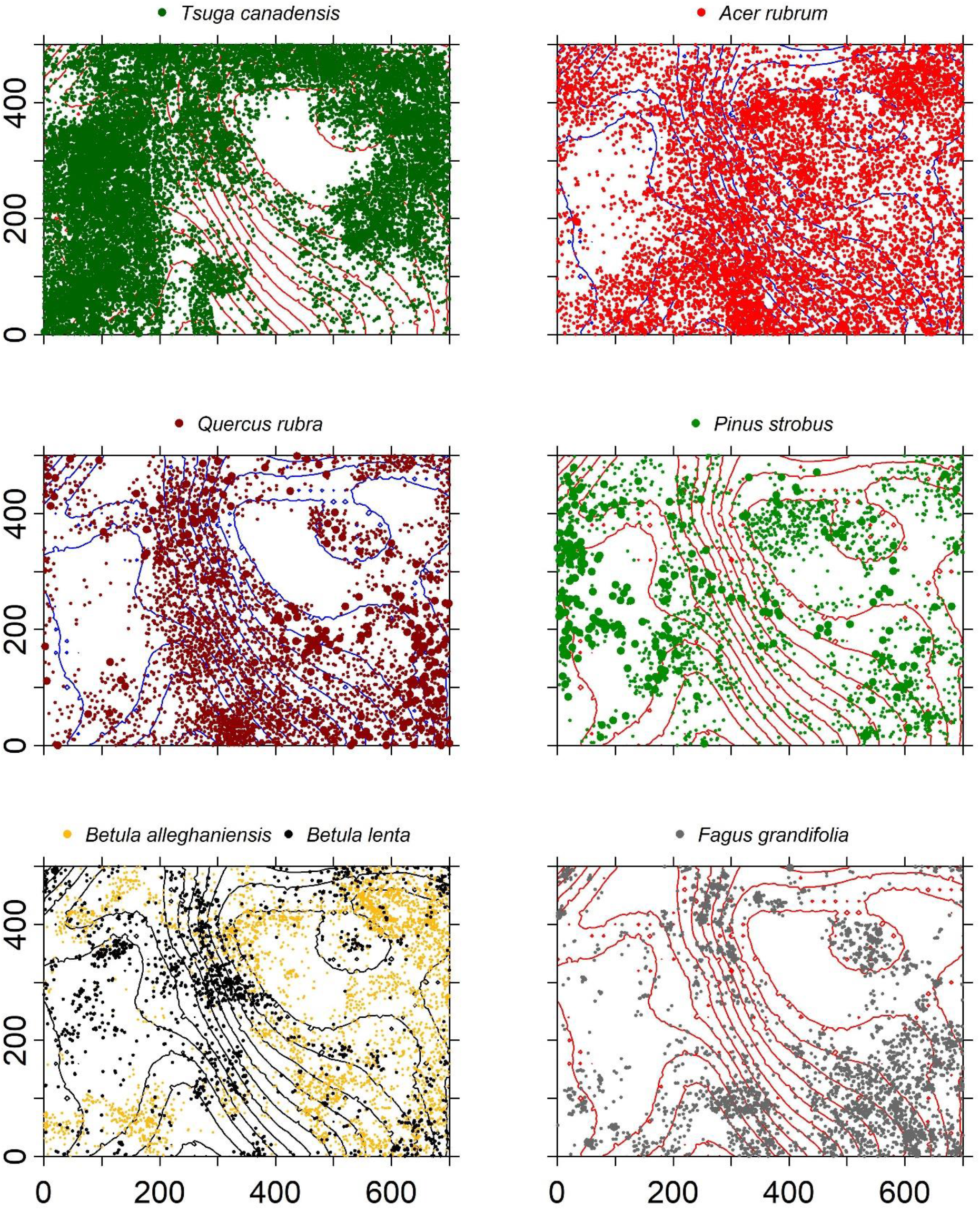
Spatial distribution of stems ≥1 cm DBH of the seven most common overstory species within the HF ForestGEO plot with 3 m elevation contour lines.

Shrubs were often found in aggregations related to hydrology and topography. *Ilex verticillata V. corymbosum*, *Viburnum nudum* (withe-rod), and *Lyonia ligustrina* (maleberry) dominated the poorly drained beaver swamp (Fig. 6). *Hamamelis virginiana* (witch-hazel) was found in a narrow elevational band (342-346 m) just above the swamp and a dense patch of *K. latifolia* was in the northwest corner of the plot.

**Figure 6.**
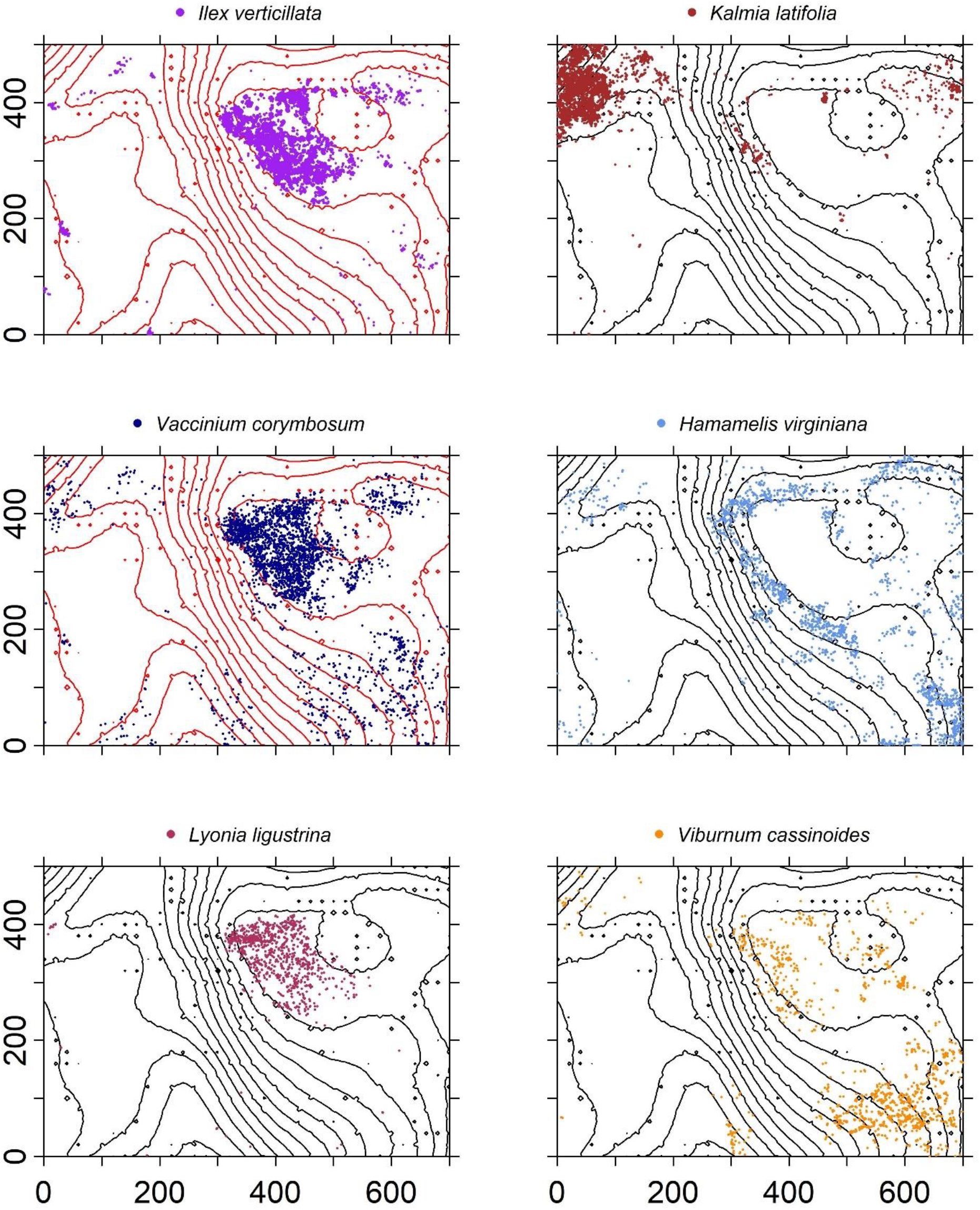
Spatial distribution of stems ≥1 cm DBH of the six most common understory species within the HF ForestGEO plot with 3 m elevation contour lines.

The seven most abundant canopy tree species were significantly clustered in the plot at all spatial lags up to 50m relative to a CSR null expectation (Fig. 7). The effect of intraspecific competition also was apparent for these seven species. Spatial distributions of *A. rubrum*, *Q. rubra*, and *F. grandifolia* showed negative intraspecific correlations in diameters up to at least a 150-m spatial lag, whereas the other species had intraspecific negative correlations at ≤ 50-m spatial lags (Fig. 8). *Tsuga canadensis*, *B. alleghaniensis*, and *P. strobus* had positive spatial correlations among DBHs at spatial lags > 150 m. Interspecific correlations in diameters between species suggest that the impact of *T. canadensis* on *Q. rubra* was negative at intermediate spatial lags (25–75 m) but positive between *T. canadensis* and the other five species at most spatial scales up to 150 m (Fig. 9).

**Figure 7.**
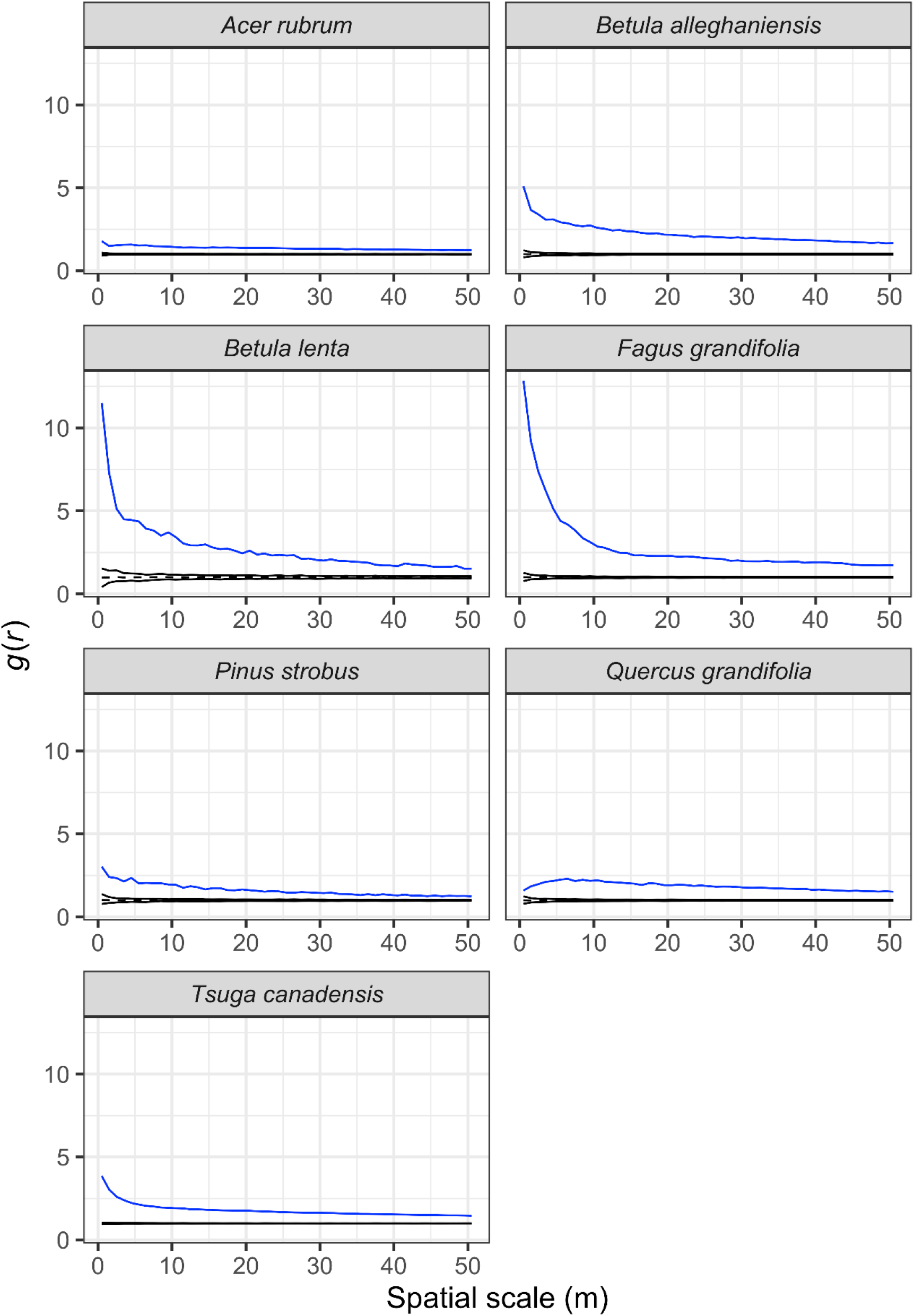
Observed (blue line) and expected (black dashed line) values of the pair correlation function, *g*(*r*), showing the degree of spatial clustering (values >1) of the seven most dominant tree species in the Harvard Forest plot. Expected values were obtained from 199 Monte Carlo simulations to completely randomize the spatial position of trees (complete spatial randomness; CSR).

**Figure 8.**
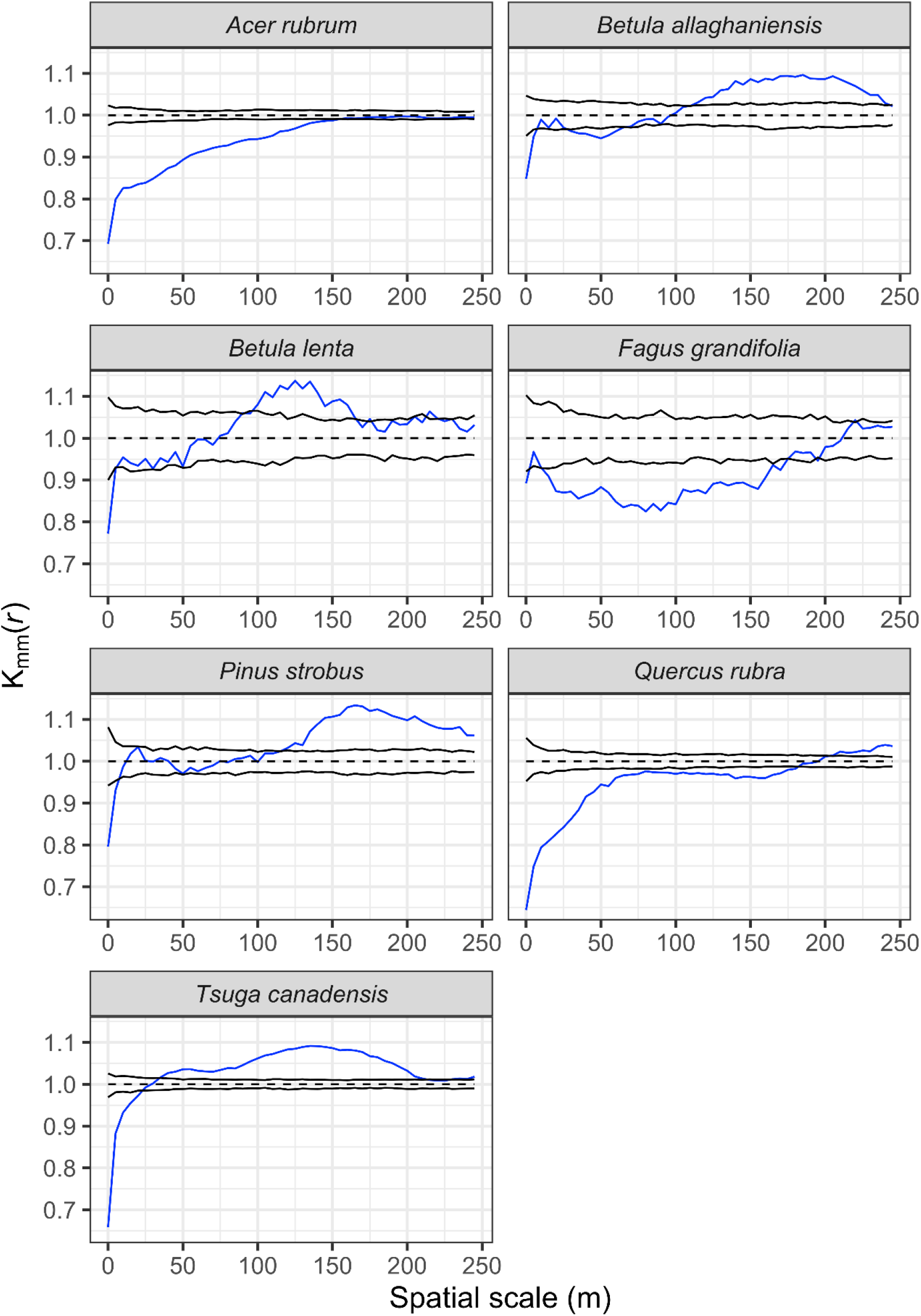
Univariate mark correlation function analysis results showing the effects of the underlying spatial pattern of trees on the size of conspecific individuals for seven dominant species in the Harvard Forest plot across a range of scales. The significance of this effect was evaluated by comparing the calculated *kmm*(*r*) against values simulated under a null expectation, where tree sizes were randomly shuffled over all trees for each of the 199 simulations. The blue line indicates calculated *kmm*(*r*) values, while the black lines demark the 95% confidence envelope around simulated *kmm*(*r*) values under the null model. A blue line falling below, within, or above the upper confidence limit, indicates significant negative, independent, or positive correlations among DBH marks for the given species, respectively.

**Figure 9.**
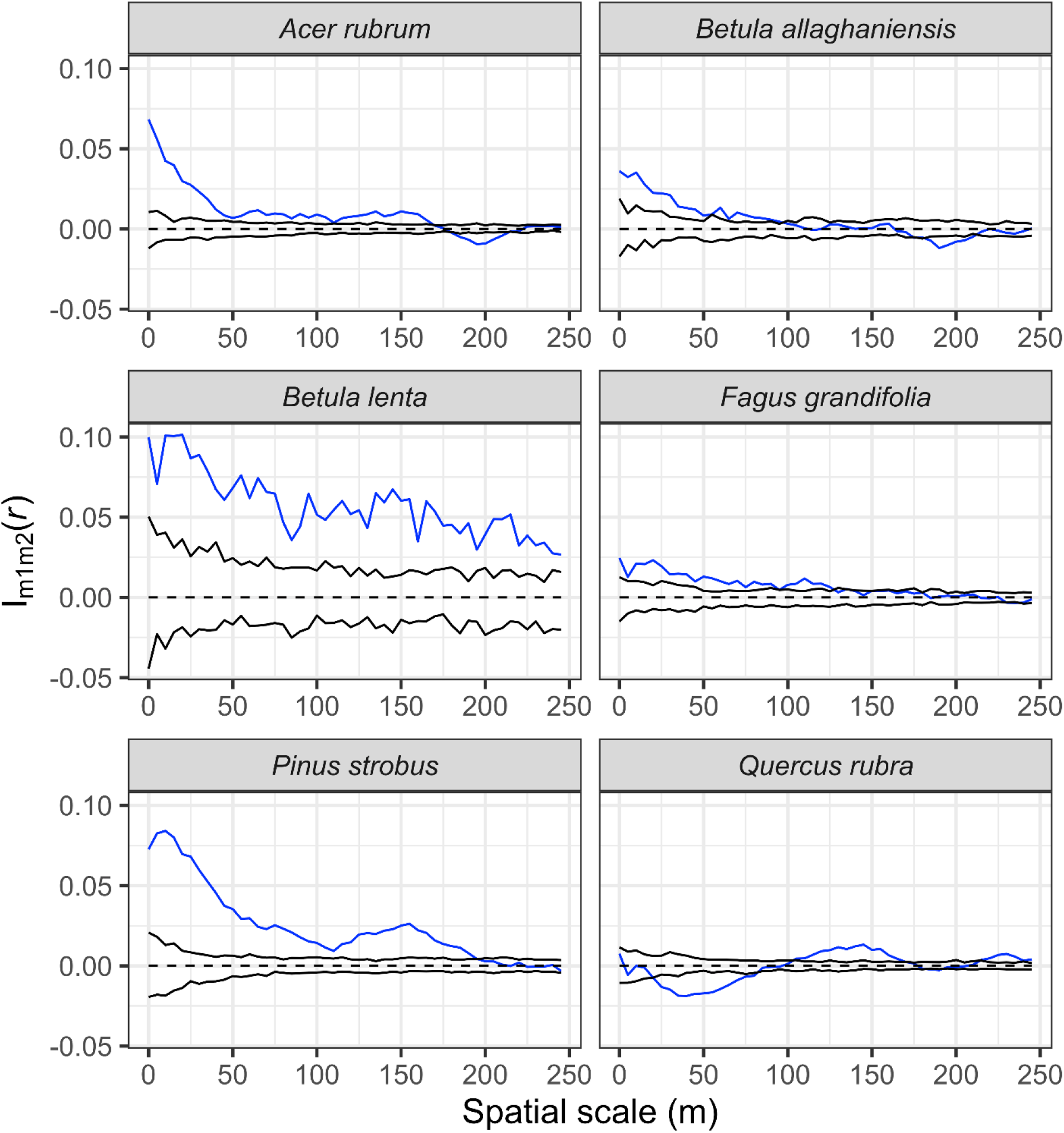
Bivariate marked point pattern analysis results showing the effects of the size of focal *Tsuga canadensis* individuals on the size of six other non-focal species in the HF ForestGEO plot across a range of scales. The significance of this effect was evaluated by comparing the calculated Schlather’s I (*Im1m2*(*r*)) bivariate correlation statistic against values simulated under a null expectation, where non-focal species’ tree sizes were randomly shuffled over trees for each of 199 simulations. The blue line indicates calculated *Im1m2*(*r*) values, while the black lines demark the 95% confidence envelope around simulated *Im1m2*(*r*) values under the null model. A blue line falling below, within, or above the upper confidence limit, indicates significant negative, independent, or positive correlations of DBH marks of the given species with the DBH of *T. canadensis* individuals found at a range of distances, respectively.

The abundances and sizes of the most common overstory species were predicted best by a variety of historical factors and competitive interactions. Conditional inference random-forest modeling revealed that the abundances of *T. canadensis*, *P. strobus, Q. rubra, A. rubrum* and *F. grandifolia* were strongly associated with neighborhood effects (size of neighboring trees within 10 m; Fig. 10). The date of field abandonment was a strong predictor of *Q. rubra*, *P. strobus*, and *A. lenta* abundance, whereas the forest type in 1908 was the best predictor of *B. alleghaniensis* and *A. rubrum* abundance. *Betula* species also were strongly associated with Simmons soil type. Overstory species diameters were best predicted by neighborhood effects for *T. canadensis*, *B. lenta*, and *F. grandifolia*; date of field abandonment for *P. strobus* and *B. alleghaniensis*; and the 1947 stand type for *Q. rubra* and *A. rubrum* (Fig. 11). The predictive power of the conditional inference forest model regressions was much higher (*R*^2^ = 0.79 – 0.95) for species abundance in the plot compared to species size (*R*^2^ = 0.11 – 0.53).

**Figure 10.**
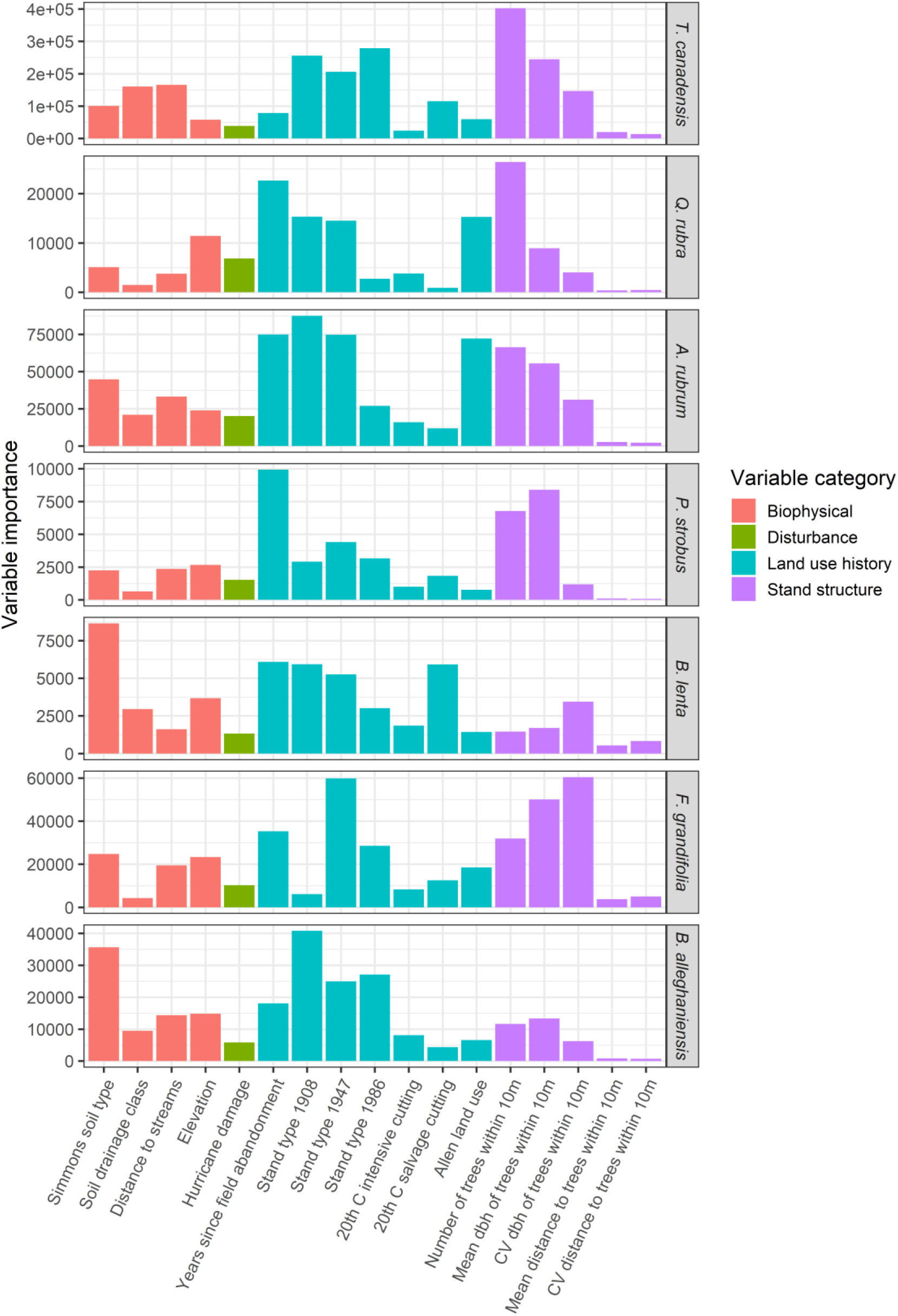
Variable importance scores, based on the mean decrease in prediction accuracy, from a conditional inference random-forest model predicting tree species abundance values (stems/ha) for the seven most common trees as a function of possible predictors (Table 1). Variable importance scores were calculated across 400 random forest iterations and the range of values is from 0-100,000, reflecting the range of the response variable, abundance.

**Figure 11.**
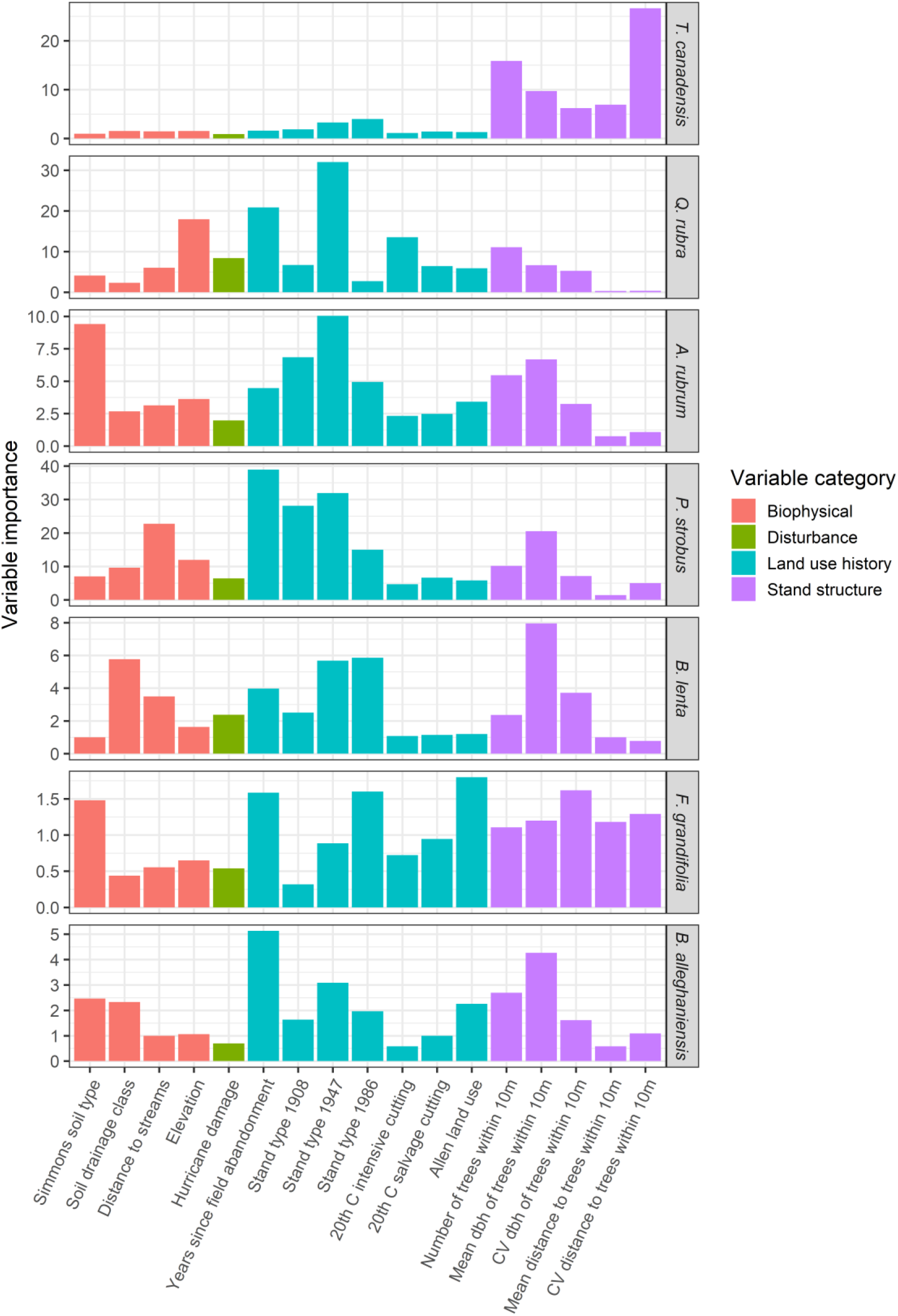
Variable importance scores, based on the mean decrease in prediction accuracy, from a conditional inference random-forest model predicting tree species diameter at breast height (DBH) for the seven most common trees as a function of possible predictors (Table 1). Variable importance scores were calculated across 400 random forest iterations and the range of values is from 0 - 40, reflecting the range of the response variable, diameter.

## Discussion

We censused all woody stems ≥ 1cm DBH within a 35-ha forest-dynamics plot in north-central Massachusetts to examine the spatial patterns of trees and shrubs at a scale rarely attempted in temperate forests. We have shown that broad patterns in land use and historical disturbance that occurred up to a century ago remain the dominant controls on present-day spatial distribution and structure of overstory species. Tree species were significantly clumped within the plot and *T. canadensis* affected the distribution of other dominant canopy species in different ways. Topography and hydrology also affected the distribution and abundance of understory stems. Detailed abundance and species distribution data provided in this study will provide invaluable information on forest dynamics in the future as the currently most abundant species—*Tsuga canadensis*—is declining because of a non-native insect (Orwig et al. 2018).

*Forest structure is contingent on past land use* The forest canopy within the HF ForestGEO plot, dominated by *T. canadensis*, *Q. rubra*, *A. rubrum*, and *P. strobus*, is representative of many central New England forests. Like other temperate ForestGEO plots, a relatively small number of species dominated the HF plot (13 species were represented by over 1000 stems). However this number was higher than the 5–10 species that reached this abundance in other temperate ForestGEO plots (Wang et al. 2010, Wang et al. 2011, Lutz et al. 2012, Bourg et al. 2013, Lutz et al. 2013) and likely reflects the varied habitats, high intensity of prior land use, and early stages of stand development at HF. Although we have much historical knowledge regarding land-use change at HF, the conditional regression random-forest modeling enabled us to explore more quantitatively how patterns of tree size and stem density for the seven most abundant species have been affected by tradeoffs between legacy effects of past land uses, management interventions, disturbances, and local-scale variation in stand structure and environmental conditions. This combination of quantitative modeling with historical knowledge contributes to a deeper understanding of historical human impacts on current forest structure.

For example, our modeling results suggested that *T. canadensis* diameters and stem densities across the full plot are most strongly associated with local stand structural characteristics and neighborhood effects, while stem densities are only moderately associated with land-use history. This result is consistent with the appearance of the 35-ha HF plot as a relatively undisturbed old forest stand, its persistence through time, and the exclusion of other species under its canopy. *T. canadensis* is most abundant on land that was consistently used as a woodlot but never completely cleared for agriculture. The western portion of the plot was one of the few locations at HF that was mapped as *T. canadensis* forest in 1908 (Spurr 1956) and where *T. canadensis* currently is most prominent. It is also the location where the presence of *Tsuga* has been documented for the last 8000 years (Foster and Zebryk 1993). The high abundance of *T. canadensis* is the result of its shade tolerance and deep crowns, which enable it to persist for decades, modify the understory environment by transmitting very little light, prevent other species from getting established (Canham et al. 1994), and gain dominance following partial cuttings, the death and subsequent salvage of *C. dentata* and *F. grandifolia*, and moderate damage from the 1938 hurricane (Foster et al. 1992, Motzkin et al. 1999, McLachlan et al. 2000). These same disturbances also likely led to growth increases and additional establishment of *P. strobus* (Hibbs 1982b); the largest pine stems also occur on the western edge of the HF plot.

In contrast, modeling revealed stronger effects of both land-use history and stand structural variables on the sizes and stem densities of the other six dominant species. Field abandonment date and stand types present in the early- and mid-20^th^ century are particularly strong predictors of diameters and densities of these species. This is consistent with recorded historical knowledge. For example, *Pinus strobus* and *Q. rubra* are most abundant on areas that were formerly pasture or fields in the mid- to late-1800s and also experienced intensive past silvicultural cuts, thinning, and weeding in the 1920s–1940s, and more severe damage from the 1938 hurricane (Motzkin et al. 1999, Hall 2005). *Quercus rubra* trees had larger mean diameters and crown sizes than *F. grandifolia* or *A. rubrum*, consistent with past investigations that highlighted the ability of *Q. rubra* to overtop canopy associates and rapidly expand laterally into gaps (Oliver 1978, Hibbs 1982a). *Acer rubrum* and *B. alleghaniensis* are more closely associated with mesic locations such as swamp borders with silt loam soils and low-lying sites with peaty soils in the northeast corner of the plot; indeed, random-forest models supported the relatively strong importance of soil type for these species and *B. lenta* relative to the other species. The south-central portion of the plot experienced the most intensive land use. It was the only area that experienced historical cultivation and multiple periods of subsequent clear-cutting, including a harvest in 1990. This area is dominated by smaller, multi-stemmed *A. rubrum*, *Q. rubra*, *B. populifolia* and *B*. *papyrifera* (grey and paper birch), and *Prunus* (cherry) species, which are much more common in forests that have experienced intense human impacts (Del Tredici 2001). The known relationships between current stem-density patterns for *A. rubrum* and the two *Betula* species and historical land-use activities are borne out by the random-forest modelling. These species sprout following cutting and take advantage of high-light environments (Burns and Honkala 1990).

Understory composition, dominated by woody shrubs, appears to be determined by soil drainage and the ability of individual species to tolerate standing water, poorly drained soils, or subtle topographic variation. Historically, the swamp contained pasture on its western edge and a woodlot in the remaining portion. Today, the wetland shrubs *I. verticillata*, *Va. corymbosum*, *L. ligustrina*, and *Vi. nudum* are found in high abundance in the central beaver swamp, which otherwise is devoid of trees. The northwest corner has the highest elevation and is dominated by *K. latifolia*. *Hamamalis virginiana* appears to be restricted to a narrow elevation west of the swamp and in the southeast corner of the plot. Previous work at HF related *K. latifolia* abundance to nitrogen-poor sites and *H. virginiana* to continuously forested sites (Motzkin et al. 1999), which is consistent with our findings.

Across all species and size classes, the forest contains a preponderance (> 80,000) of small stems (< 10-cm DBH) that exhibit a reverse-J size distribution. The high abundance of stems in this size class (e.g., several shrub species, *T. canadensis,* and *A. rubrum*) is in contrast to several other temperate forest plots (Lutz et al. 2012, Bourg et al. 2013, Lutz et al. 2013), and is more similar to results from tropical evergreen (Memiaghe et al. 2016) or Mediterranean forests (Gilbert et al. 2010). Most of the abundant overstory and all the abundant shrub species also have reverse-J distributions, indicative of stable populations and adequate regeneration. For overstory species, this likely is a result of a mix of even-age and varying-aged cohorts and single trees establishing following anthropogenic disturbances and natural gap-phase dynamics that are frequent in this region (Oliver and Stephens 1977, Hibbs 1982a, Pederson 2005). The greater ages of the shade-tolerant *T. canadensis* that occur on primary woodland are approaching a structure and diameter distribution that resembles old-growth forest (D’Amato et al. 2008, Janowiak et al. 2008). In contrast, *A. rubrum* and *Q. rubra* had skewed unimodal size distributions more indicative of managed forests (Janowiak et al. 2008).

### Overstory spatial patterns

We observed significant spatial clustering among abundant overstory species at all spatial scales examined. Aggregated species distribution patterns are common in both temperate (Hou et al. 2004, Hao et al. 2007, Wang et al. 2011) and tropical forests (Condit et al. 2000, Plotkin et al. 2000, Réjou-Méchain et al. 2011, Nguyen et al. 2016). Both external factors (habitat heterogeneity) and internal factors (dispersal limitation, succession, gap dynamics) can lead to clumped distributions at various spatial scales (Getzin et al. 2008, Réjou-Méchain et al. 2011).

Within the HF ForestGEO plot, high habitat heterogeneity caused by complex past land use (Motzkin et al. 1999) has led to high densities of *A. rubrum* and *Q. rubra* stems in the central portion where the most intensive land use occurred in the past. These non-random patches of individuals with lower than average DBH (as seen in the mark correlation analysis) may reflect strong competition for light as seen elsewhere (Fibich et al. 2016). Similar patterns seen in *B. alleghaniensis*, *B*. *lenta*, *P. strobus*, and *T. canadensis* in close proximity to other conspecifics (0–20-m scale) likely reflect crowding effects, and for *T. canadensis*, the ability of thousands of small stems to persist in the understory for decades (Marshall 1927). These effects disappear at intermediate scales and even become positive at distances > 100 m, indicating that trees greater than the mean DBH are more broadly distributed. The negative correlation observed for *F. grandifolia* at most spatial lags ≤ 150 m may be more reflective of its overall size distribution with most of its stems < 10 cm DBH. Beech-bark disease is present at HF, and has likely contributed, along with past cutting, to the absence of large *F. grandifolia* in the plot.

Bivariate mark correlation functions have been underused in large, stem-mapped plots but hold great promise in ecological research (Velázquez et al. 2016). We used this method to examine the relationship between the size of individuals of *T. canadensis*, an important foundation species within the plot, with the size of six other important canopy species some distance away. Apart from *Q. rubra*, diameters of the other five species were positively correlated with the diameters of *T. canadensis* at all spatial scales. This pattern is consistent with *T. canadensis* being a foundation species in this forest (Buckley et al. 2016, Ellison et al. 2019), but it also simply could indicate a “habitat” effect: all these species are growing well everywhere and are found at a wide range of sizes. This effect was particularly strong for *B. lenta* and *P. strobus*. This effect was weaker for *A. rubrum*, *B. alleghaniensis*, and *F. grandifolia* and disappeared after 100–150 m. Diameter of *Q. rubra* was on average smaller than expected by chance when within 20–80 m of *T. canadensis*. Historical factors play a role here, as the spatial distribution of these species highlight that oak abundance is the lowest within the *T. canadensis*-dominated portions of the plot that were woodlots and suggest that *T. canadensis* and the dense shade cast by their crowns limited establishment of the more intolerant *Q. rubra*.

### Summary

The HF ForestGEO plot is the largest mapped temperate-forest plot in North America and joins the growing array of temperate plots worldwide (Anderson-Teixeira et al. 2015). The species composition and aggregated spatial patterns within the plot are still being influenced by anthropogenic and natural disturbances that occurred decades to over a century ago. Despite extensive 20^th^-century harvesting, silvicultural thinning, and salvage operations following the 1938 hurricane, the most common overstory species in the HF ForestGEO plot today can best be predicted by longer-term land-use legacies represented by the 1908 forest type and the date of late 19^th^-century field abandonment, and tree neighborhood effects. At smaller scales, there is evidence of crowding effects of many common species, likely due to successional dynamics of these aggrading forests following intensive land use. The increasing importance of *T. canadensis* during the last century across the plot negatively affected the distribution of *Q. rubra*. Its location and five-year schedule of plot sampling highlight the plot as valuable long-term infrastructure that will complement Harvard Forest, LTER, NEON, and ForestGEO research efforts (Orwig et al. 2018). Because all woody stems ≥ 1 cm DBH are mapped and measured, the data have been used in a variety of complementary ways including to examine species codispersion patterns and spatial patterns of species co-occurrence (Buckley et al. 2016, Case et al. 2016), help inform a simulation model of forest dynamics (SORTIE (Case et al. 2017)), assist with investigating crown allometry (Sullivan et al. 2017) and mapping (Hastings et al. 2020), and aid in identifying statistical fingerprints of foundation species (Ellison et al. 2019). In addition, the data enable us to document changing species distribution patterns at an uncommonly large scale, while focusing on elements of the landscape that are often ignored, like beaver swamps and shrub thickets, and examine their contribution to overall forest structure, composition, and related hydrology.

## Acknowledgements

Funding for the Harvard ForestGEO Forest Dynamics plot was provided by the Center for Tropical Forest Science and Smithsonian Institute’s Forest Global Earth Observatory (CTFS-ForestGEO), the National Science Foundation’s LTER program (DEB 06-20443, DEB 12-37491, DEB 18-32210) and Harvard University. We thank the many field technicians who helped census the plot including Jerry Breault, Kyle Gay, Mike Babineau, Christian Foster, Jeff Hutchins, Tamara Martz, Elizabeth Barnes, Rachel DeMatte, Bianca Kubierschky, Ellen Kujawa, Audrey Lamb, Ben Misiuk, Finn Olcott, Hallie Schwab, Alida Mau, Joe Horn, Taylor Lucey, Brett Gelinas, Danelle Laflower, Sarah Meyers, and Kyle Krigest. We are grateful to John Wisnewski and the woods crew at HF for providing materials, supplies, and invaluable field assistance with plot logistics. Joel Botti and Frank Schiappa provided survey expertise to establish the 35-ha plot. Special thanks to Stuart Davies and Rick Condit for field training, database assistance, and plot advice. Sean McMahon and Suzanne Lao were extremely helpful with field planning, data questions, and many plot logistics. Thanks to Jeannette Bowlen for administrative assistance, Emery Boose and Paul Siqueira for help with plot coordinates, Brian Hall for map and GIS layer production, and Matthew Duveneck and Danelle Laflower for assistance with figures. We also thank David Foster for his support and assistance with plot design, location, and integration with other long-term studies at HF. Jonathan Thompson, Neil Pederson, Audrey Barker-Plotkin and lab group members provided critical comments on earlier versions of the manuscript. There were no conflicts of interest.

---

**Table S1.**
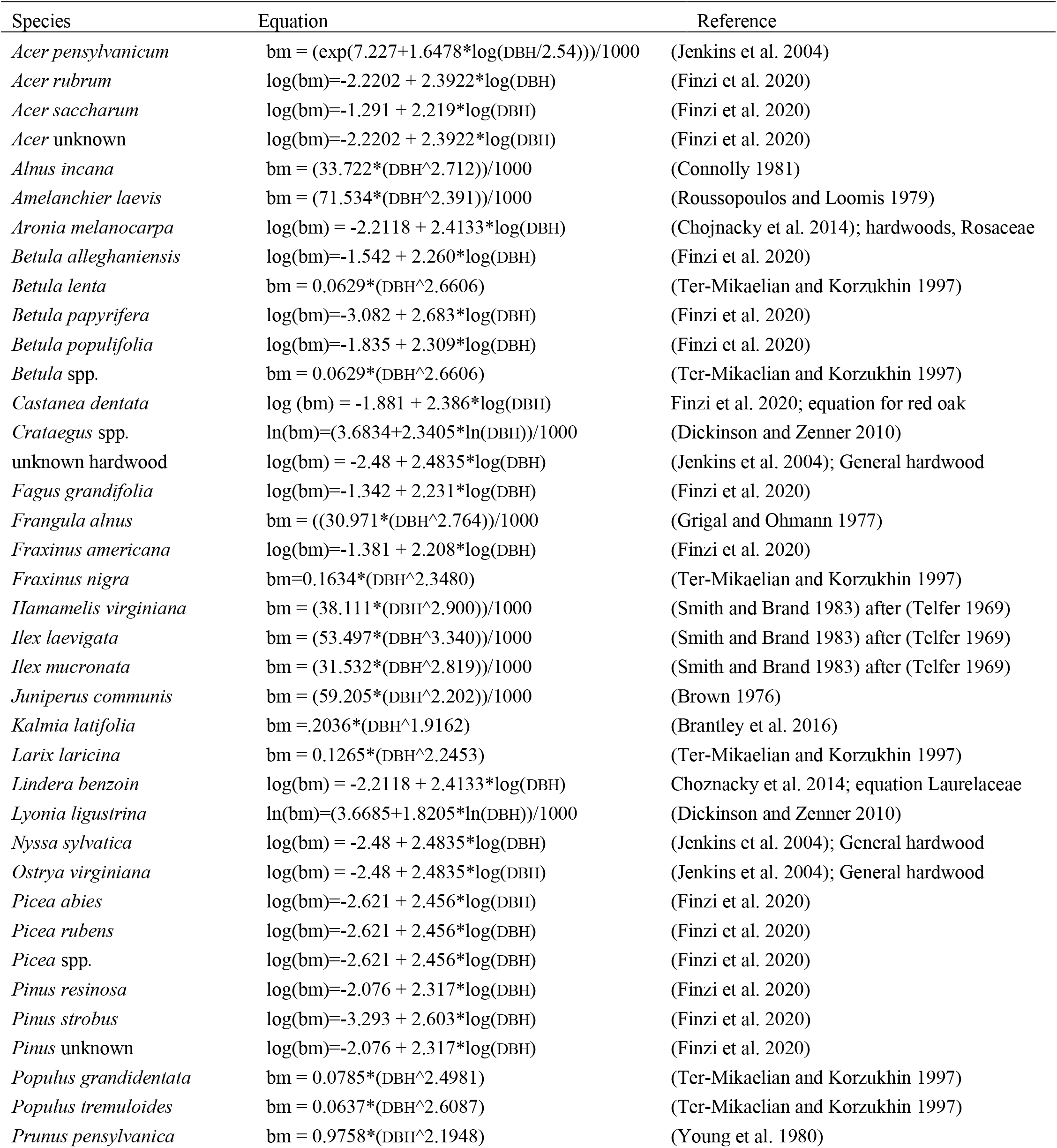
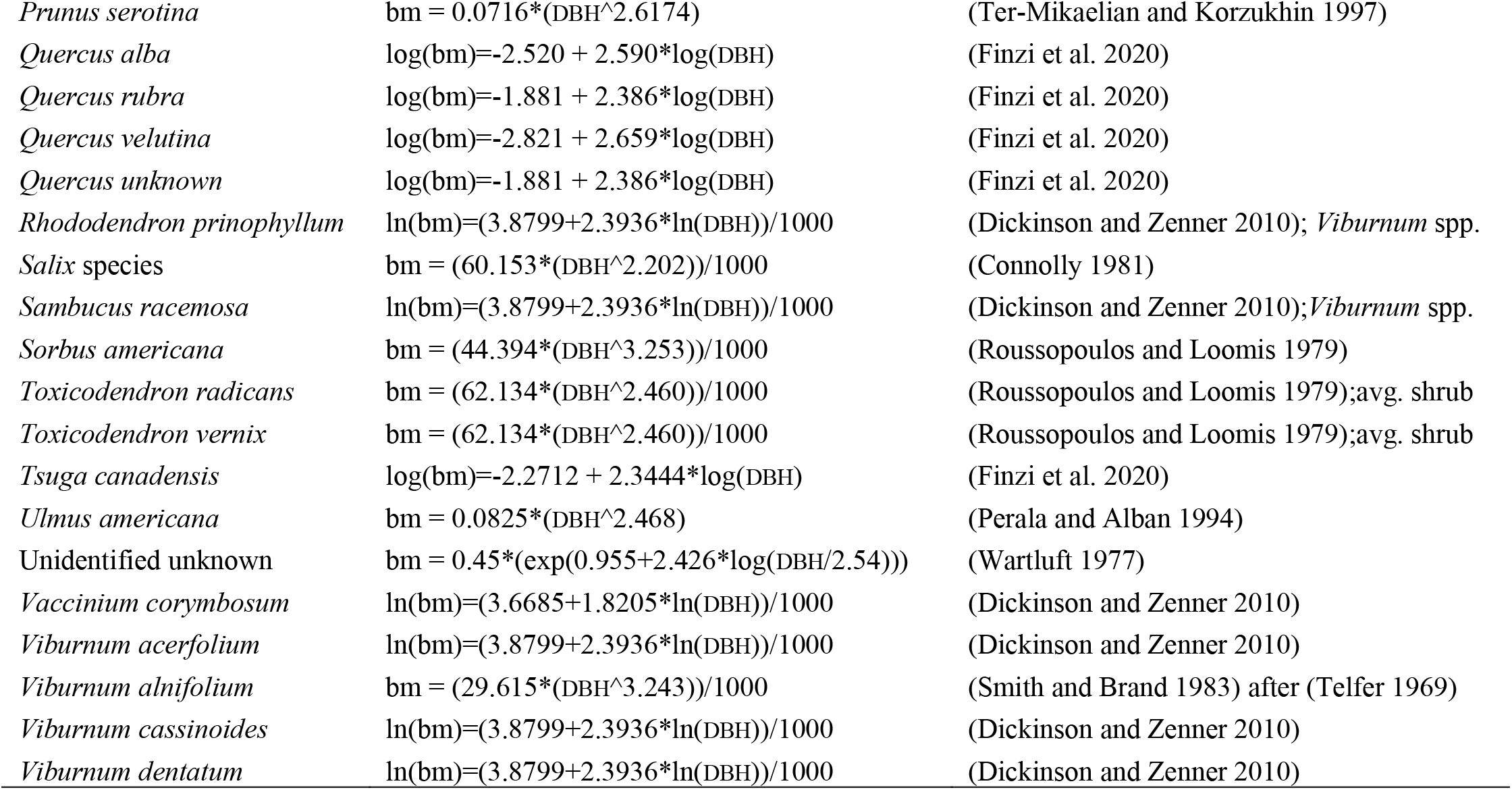
Biomass equations of woody species within the HF ForestGEO plot. bm = biomass (kg), DBH = diameter at breast height (cm).

**Table S2.**
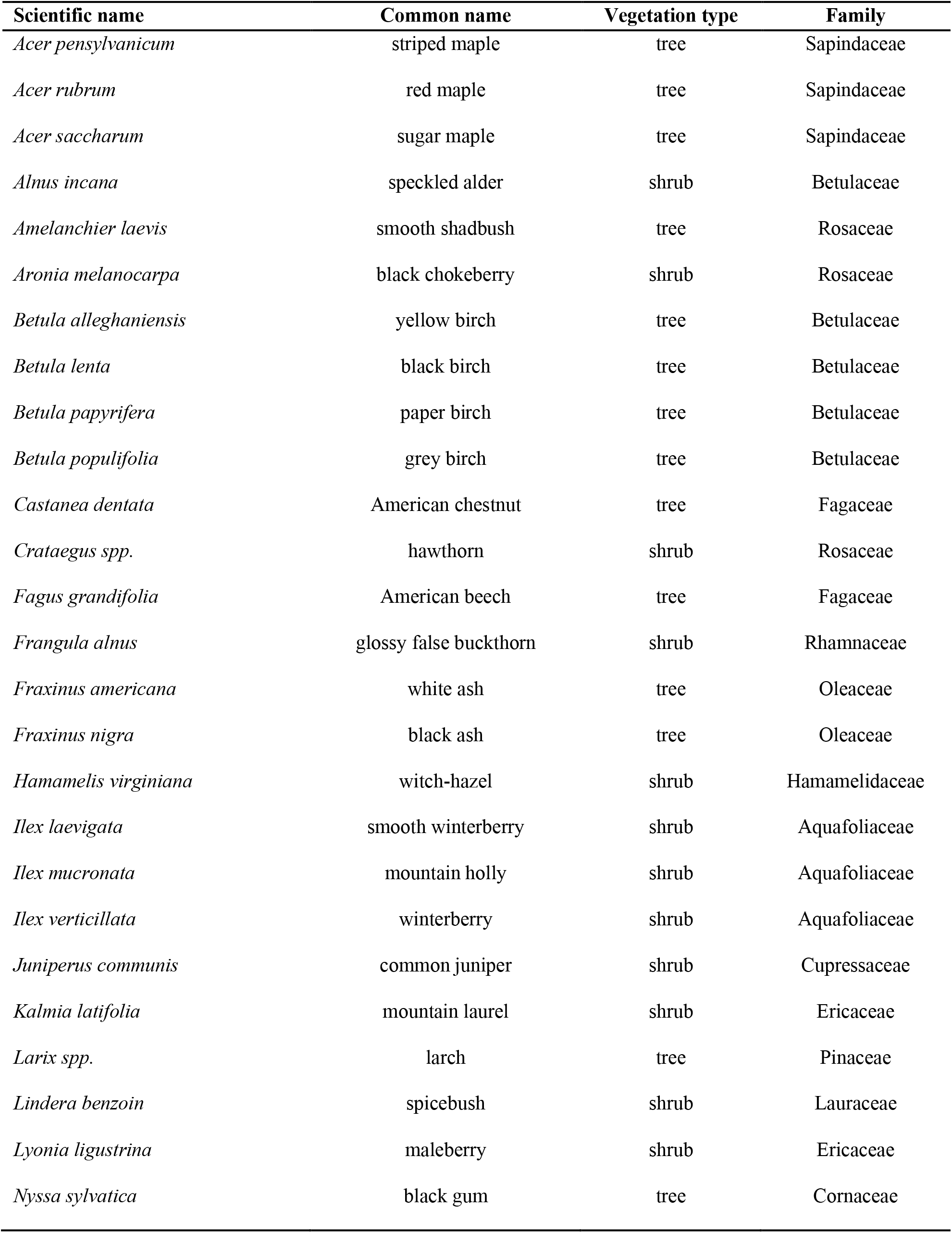
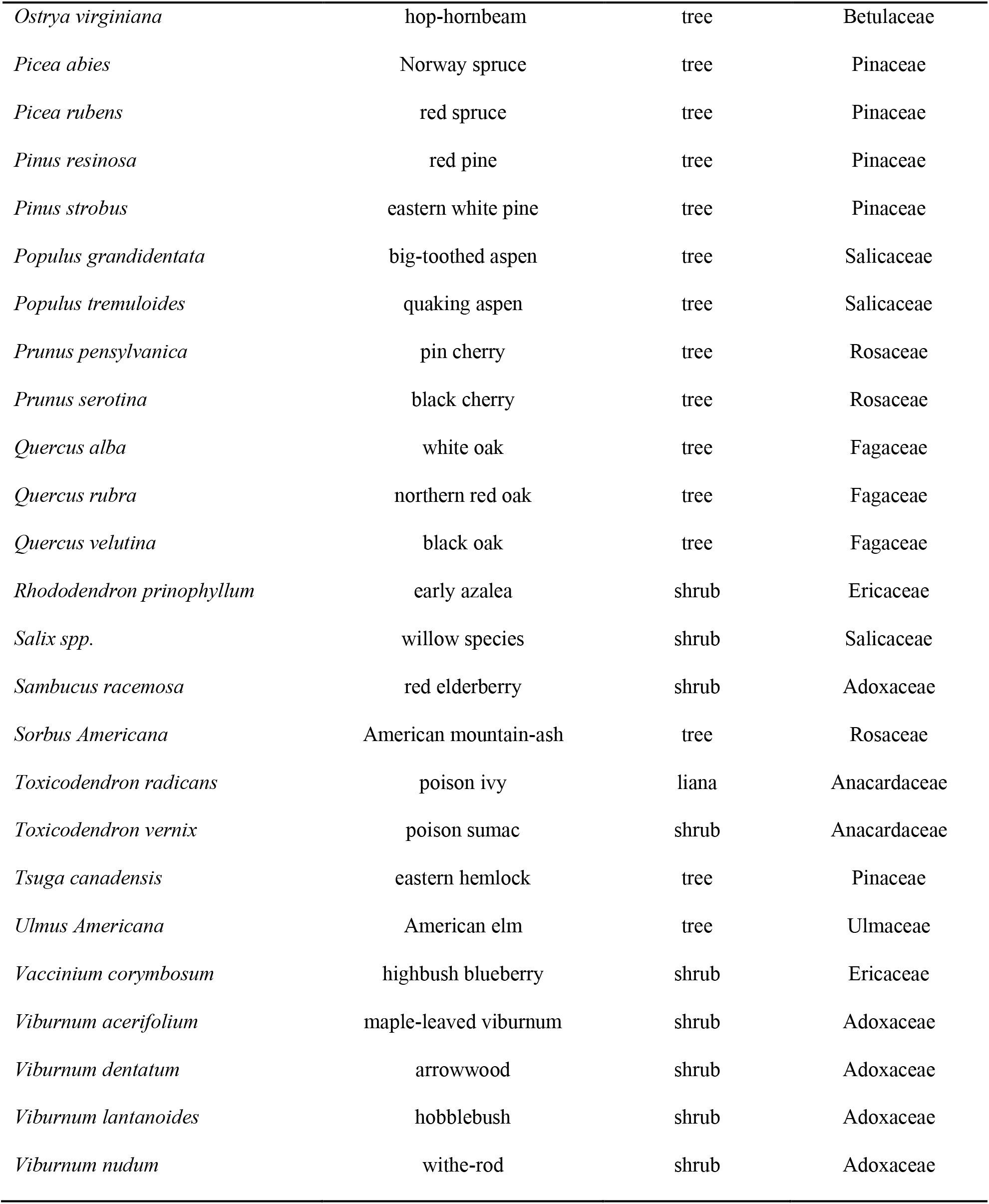
List of woody plant species ≥ 1 cm DBH within the HF ForestGEO plot in 2014.

| Scientific name | Common name | Vegetation type | Family |
| --- | --- | --- | --- |
| *Acer pensylvanicum* | striped maple | tree | Sapindaceae |
| *Acer rubrum* | red maple | tree | Sapindaceae |
| *Acer saccharum* | sugar maple | tree | Sapindaceae |
| *Alnus incana* | speckled alder | shrub | Betulaceae |
| *Amelanchier laevis* | smooth shadbush | tree | Rosaceae |
| *Aronia melanocarpa* | black chokeberry | shrub | Rosaceae |
| *Betula alleghaniensis* | yellow birch | tree | Betulaceae |
| *Betula lenta* | black birch | tree | Betulaceae |
| *Betula papyrifera* | paper birch | tree | Betulaceae |
| *Betula populifolia* | grey birch | tree | Betulaceae |
| *Castanea dentata* | American chestnut | tree | Fagaceae |
| *Crataegus spp.* | hawthorn | shrub | Rosaceae |
| *Fagus grandifolia* | American beech | tree | Fagaceae |
| *Frangula alnus* | glossy false buckthorn | shrub | Rhamnaceae |
| *Fraxinus americana* | white ash | tree | Oleaceae |
| *Fraxinus nigra* | black ash | tree | Oleaceae |
| *Hamamelis virginiana* | witch-hazel | shrub | Hamamelidaceae |
| *Ilex laevigata* | smooth winterberry | shrub | Aquafoliaceae |
| *Ilex mucronata* | mountain holly | shrub | Aquafoliaceae |
| *Ilex verticillata* | winterberry | shrub | Aquafoliaceae |
| *Juniperus communis* | common juniper | shrub | Cupressaceae |
| *Kalmia latifolia* | mountain laurel | shrub | Ericaceae |
| *Larix spp.* | larch | tree | Pinaceae |
| *Lindera benzoin* | spicebush | shrub | Lauraceae |
| *Lyonia ligustrina* | maleberry | shrub | Ericaceae |
| *Nyssa sylvatica* | black gum | tree | Cornaceae |
| *Ostrya virginiana* | hop-hornbeam | tree | Betulaceae |
| *Picea abies* | Norway spruce | tree | Pinaceae |
| *Picea rubens* | red spruce | tree | Pinaceae |
| *Pinus resinosa* | red pine | tree | Pinaceae |
| *Pinus strobus* | eastern white pine | tree | Pinaceae |
| *Populus grandidentata* | big-toothed aspen | tree | Salicaceae |
| *Populus tremuloides* | quaking aspen | tree | Salicaceae |
| *Prunus pensylvanica* | pin cherry | tree | Rosaceae |
| *Prunus serotina* | black cherry | tree | Rosaceae |
| *Quercus alba* | white oak | tree | Fagaceae |
| *Quercus rubra* | northern red oak | tree | Fagaceae |
| *Quercus velutina* | black oak | tree | Fagaceae |
| *Rhododendron prinophyllum* | early azalea | shrub | Ericaceae |
| *Salix spp.* | willow species | shrub | Salicaceae |
| *Sambucus racemosa* | red elderberry | shrub | Adoxaceae |
| *Sorbus Americana* | American mountain-ash | tree | Rosaceae |
| *Toxicodendron radicans* | poison ivy | liana | Anacardaceae |
| *Toxicodendron vernix* | poison sumac | shrub | Anacardaceae |
| *Tsuga canadensis* | eastern hemlock | tree | Pinaceae |
| *Ulmus Americana* | American elm | tree | Ulmaceae |
| *Vaccinium corymbosum* | highbush blueberry | shrub | Ericaceae |
| *Viburnum acerifolium* | maple-leaved viburnum | shrub | Adoxaceae |
| *Viburnum dentatum* | arrowwood | shrub | Adoxaceae |
| *Viburnum lantanoides* | hobblebush | shrub | Adoxaceae |
| *Viburnum nudum* | withe-rod | shrub | Adoxaceae |

## Literature cited

Anderson-Teixeira, K. J., S. J. Davies, A. C. Bennett, E. B. Gonzalez-Akre, H. C. Muller-Landau, S. Joseph Wright, K. Abu Salim, A. M. Almeyda Zambrano, A. Alonso, and J. L. Baltzer. 2015. CTFS-Forest GEO: a worldwide network monitoring forests in an era of global change. Global Change Biology 21:528–549.

Birks, H. H., H. Birks, P. E. Kaland, and D. Moe. 1988. The cultural landscape: past, present and future. Cambridge University Press.

Boose, E., and E. Gould. 2019. Harvard Forest Climate Data since 1964. Harvard Forest Data Archive: HF300. available online: http://harvardforest.fas.harvard.edu.

Bourg, N. A., W. J. McShea, J. R. Thompson, J. C. McGarvey, and X. Shen. 2013. Initial census, woody seedling, seed rain, and stand structure data for the SCBI SIGEO Large Forest Dynamics Plot. Ecology 94:2111–2112.

Brantley, S. T., M. L. Schulte, P. V. Bolstad, and C. F. Miniat. 2016. Equations for estimating biomass, foliage area, and sapwood of small trees in the southern Appalachians. Forest Science 62:414–421.

Breiman, L. 2001. Random forests. Machine learning 45:5–32.

Brown, J. K. 1976. Estimating shrub biomass from basal stem diameters. Canadian Journal of Forest Research 6:153–158.

Buckley, H. L., B. S. Case, and A. M. Ellison. 2016. Using codispersion analysis to characterize spatial patterns in species co-occurrences. Ecology 97:32–39.

Burns, R. M., and B. H. Honkala. 1990. Silvics of North America. Volume 2. Hardwoods. United States Department of Agriculture Washington, DC.

Canham, C. D., A. C. Finzi, S. W. Pacala, and D. H. Burbank. 1994. Causes and consequences of resource heterogeneity in forests: interspecific variation in light transmission by canopy trees. Canadian Journal of Forest Research 24:337–349.

Case, B. S., H. L. Buckley, A. A. Barker-Plotkin, D. A. Orwig, and A. M. Ellison. 2017. When a foundation crumbles: forecasting forest dynamics following the decline of the foundation species Tsuga canadensis. Ecosphere 8:e01893.

Case, B. S., H. L. Buckley, A. Barker Plotkin, and A. Ellison. 2016. Using codispersion analysis to quantify temporal changes in the spatial pattern of forest stand structure. Journal of Chilean Statistics 7:3–15.

Chazdon, R. L. 2003. Tropical forest recovery: legacies of human impact and natural disturbances. Perspectives in Plant Ecology, evolution and systematics 6:51–71.

Chojnacky, D. C., L. S. Heath, and J. C. Jenkins. 2014. Updated generalized biomass equations for North American tree species. Forestry 87:129–151.

Condit, R. 1998. Tropical forest census plots: methods and results from Barro Colorado Island, Panama and a comparison with other plots. Springer Science & Business Media.

Condit, R. 2014. CTFS R Package. Smithsonian Tropical Research Institute.

Condit, R., P. S. Ashton, P. Baker, S. Bunyavejchewin, S. Gunatilleke, N. Gunatilleke, S. P. Hubbell, R. B. Foster, A. Itoh, and J. V. LaFrankie. 2000. Spatial patterns in the distribution of tropical tree species. Science 288:1414–1418.

Connolly, B. J. 1981. Shrub Biomass--soil Relationships in Minnesota Wetlands. University of Minnesota.

D’Amato, A. W., D. A. Orwig, and D. R. Foster. 2008. The influence of successional processes and disturbance on the structure of Tsuga canadensis forests. Ecological Applications 18:1182–1199.

Del Tredici, P. 2001. Sprouting in temperate trees: a morphological and ecological review. The botanical review 67:121–140.

Dickinson, Y. L., and E. K. Zenner. 2010. Allometric equations for the aboveground biomass of selected common eastern hardwood understory species. Northern Journal of Applied Forestry 27:160–165.

Ellison, A. M. 2019. Foundation species, non-trophic interactions, and the value of being common. Iscience 13:254–268.

Ellison, A. M., M. S. Bank, B. D. Clinton, E. A. Colburn, K. Elliott, C. R. Ford, D. R. Foster, B. D. Kloeppel, J. D. Knoepp, and G. M. Lovett. 2005. Loss of foundation species: consequences for the structure and dynamics of forested ecosystems. Frontiers in Ecology and the Environment 3:479–486.

Ellison, A. M., H. L. Buckley, B. S. Case, D. Cardenas, Á. J. Duque, J. A. Lutz, J. A. Myers, D. A. Orwig, and J. K. Zimmerman. 2019. Species Diversity Associated with Foundation Species in Temperate and Tropical Forests. Forests 10:128.

Ellison, A. M., M. Lavine, P. B. Kerson, A. A. Barker Plotkin, and D. A. Orwig. 2014. Building a foundation: Land-use history and dendrochronology reveal temporal dynamics of a Tsuga canadensis (Pinaceae) forest. Rhodora 116:377–427.

Fibich, P., J. Lepš, V. Novotný, P. Klimeš, J. Těšitel, K. Molem, K. Damas, and G. D. Weiblen. 2016. Spatial patterns of tree species distribution in New Guinea primary and secondary lowland rain forest. Journal of Vegetation Science 27:328–339.

Finzi, A. C., M.-A. Giasson, A. A. Barker Plotkin, J. D. Aber, E. R. Boose, E. A. Davidson, M. C. Dietze, A. M. Ellison, S. D. Frey, E. Goldman, T. F. Keenan, J. M. Melillo, J. W. Munger, K. J. Nadelhoffer, S. V. Ollinger, D. A. Orwig, N. Pederson, A. D. Richardson, K. Savage, J. Tang, J. R. Thompson, C. A. Williams, S. C. Wofsy, Z. Zhou, and D. R. Foster. 2020. Carbon budget of the Harvard Forest Long-Term Ecological Research site: pattern, process, and response to global change. Ecological Monographs In Press.

Fisher, R. T. 1933. New England forests: biological factors. New England’s Prospect. American Geographical Society. Special Publication:213–223.

Foster, D. 2014. Hemlock: a forest giant on the edge. Yale University Press, New Haven, Connecticut, USA.

Foster, D. R. 1992. Land-use history (1730-1990) and vegetation dynamics in central New England, USA. Journal of ecology:753–771.

Foster, D. R., and J. Aber. 2004. Forests in Time. Ecosystem Structure and Function as a Consequence of 1000 Years of Change. Synthesis volume of the Harvard Forest LTER Program. Yale University Press, New Haven, Connecticut, USA.

Foster, D. R., and E. R. Boose. 1992. Patterns of forest damage resulting from catastrophic wind in central New England, USA. Journal of ecology:79–98.

Foster, D. R., G. Motzkin, and B. Slater. 1998. Land-use history as long-term broad-scale disturbance: regional forest dynamics in central New England. Ecosystems 1:96–119.

Foster, D. R., T. Zebryk, P. Schoonmaker, and A. Lezberg. 1992. Post-settlement history of human land-use and vegetation dynamics of a Tsuga canadensis (hemlock) woodlot in central New England. Journal of ecology:773–786.

Foster, D. R., and T. M. Zebryk. 1993. Long-term vegetation dynamics and disturbance history of a Tsuga-dominated forest in New England. Ecology 74:982–998.

Fox, E. W., R. A. Hill, S. G. Leibowitz, A. R. Olsen, D. J. Thornbrugh, and M. H. Weber. 2017. Assessing the accuracy and stability of variable selection methods for random forest modeling in ecology. Environmental monitoring and assessment 189:1–20.

Getzin, S., T. Wiegand, K. Wiegand, and F. He. 2008. Heterogeneity influences spatial patterns and demographics in forest stands. Journal of ecology 96:807–820.

Gilbert, G. S., E. Howard, B. Ayala-Orozco, M. Bonilla-Moheno, J. Cummings, S. Langridge, I. M. Parker, J. Pasari, D. Schweizer, and S. Swope. 2010. Beyond the tropics: forest structure in a temperate forest mapped plot. Journal of Vegetation Science 21:388–405.

Glitzenstein, J. S., C. D. Canham, M. J. McDonnell, and D. R. Streng. 1990. Effects of environment and land-use history on upland forests of the Cary Arboretum, Hudson Valley, New York. Bulletin of the Torrey Botanical Club:106–122.

Griffith, G., J. Omernik, S. Pierson, and C. Kiilsgaard. 1994. The Massachusetts Ecological Regions Project. US Environmental Protection Agency. Environmental Research Laboratory, Corvallis, OR, USA.

Grigal, D. F., and L. F. Ohmann. 1977. Biomass estimation for some shrubs from northeastern Minnesota.

Haines, A. 2011. New England Wild Flower Society’s Flora Novae Angliae: a manual for the identification of native and naturalized higher vascular plants of New England. Yale University Press.

Hall, B. 2005. Harvard Forest Properties GIS. Harvard Forest Data Archive, HF110.

Hall, B., G. Motzkin, D. R. Foster, M. Syfert, and J. Burk. 2002. Three hundred years of forest and land-use change in Massachusetts, USA. Journal of Biogeography 29:1319–1335.

Hao, Z., J. Zhang, B. Song, J. Ye, and B. Li. 2007. Vertical structure and spatial associations of dominant tree species in an old-growth temperate forest. Forest Ecology and Management 252:1–11.

Hastings, J. H., S. V. Ollinger, A. P. Ouimette, R. Sanders-DeMott, M. W. Palace, M. J. Ducey, F. B. Sullivan, D. Basler, and D. A. Orwig. 2020. Tree Species Traits Determine the Success of LiDAR-Based Crown Mapping in a Mixed Temperate Forest. Remote Sensing 12:309.

Hibbs, D. E. 1982a. Gap dynamics in a hemlock–hardwood forest. Canadian Journal of Forest Research 12:522–527.

Hibbs, D. E. 1982b. White pine in the transition hardwood forest. Canadian Journal of Botany 60:2046–2053.

Hogan, J. A., J. K. Zimmerman, J. Thompson, C. J. Nytch, and M. Uriarte. 2016. The interaction of land-use legacies and hurricane disturbance in subtropical wet forest: twenty-one years of change. Ecosphere 7:e01405.

Hothorn, T., K. Hornik, C. Strobl, A. Zeileis, and M. T. Hothorn. 2013. Package ‘party’. Citeseer.

Hothorn, T., K. Hornik, and A. Zeileis. 2006. Unbiased recursive partitioning: A conditional inference framework. Journal of Computational and Graphical statistics 15:651–674.

Hou, J., X. Mi, C. Liu, and K. Ma. 2004. Spatial patterns and associations in a Quercus-Betula forest in northern China. Journal of Vegetation Science 15:407–414.

Jacquemyn, H., P. Endels, O. Honnay, and T. Wiegand. 2010. Evaluating management interventions in small populations of a perennial herb Primula vulgaris using spatio-temporal analyses of point patterns. Journal of Applied Ecology 47:431–440.

Janowiak, M. K., L. M. Nagel, and C. R. Webster. 2008. Spatial scale and stand structure in northern hardwood forests: implications for quantifying diameter distributions. Forest Science 54:497–506.

Jenkins, J. C., D. C. Chojnacky, L. S. Heath, and R. A. Birdsey. 2004. Comprehensive database of diameter-based biomass regressions for North American tree species. Gen. Tech. Rep. NE-319. Newtown Square, PA: US Department of Agriculture, Forest Service, Northeastern Research Station. 45 p.[1 CD-ROM]. 319.

Lutz, J. A., A. J. Larson, J. A. Freund, M. E. Swanson, and K. J. Bible. 2013. The importance of large-diameter trees to forest structural heterogeneity. PLOS ONE 8:e82784.

Lutz, J. A., A. J. Larson, M. E. Swanson, and J. A. Freund. 2012. Ecological importance of large-diameter trees in a temperate mixed-conifer forest. PLOS ONE 7:e36131.

Marshall, R. 1927. The growth of hemlock before and after release from suppression. Harvard Forest.

McLachlan, J. S., D. R. Foster, and F. Menalled. 2000. Anthropogenic ties to late-successional structure and composition in four New England hemlock stands. Ecology 81:717–733.

Memiaghe, H. R., J. A. Lutz, L. Korte, A. Alonso, and D. Kenfack. 2016. Ecological importance of small-diameter trees to the structure, diversity and biomass of a tropical evergreen forest at Rabi, Gabon. PLOS ONE 11:e0154988.

Mi, C., F. Huettmann, Y. Guo, X. Han, and L. Wen. 2017. Why choose Random Forest to predict rare species distribution with few samples in large undersampled areas? Three Asian crane species models provide supporting evidence. PeerJ 5:e2849.

Mohapatra, J., C. P. Singh, M. Hamid, A. Verma, S. C. Semwal, B. Gajmer, A. A. Khuroo, A. Kumar, M. C. Nautiyal, and N. Sharma. 2019. Modelling Betula utilis distribution in response to climate-warming scenarios in Hindu-Kush Himalaya using random forest. Biodiversity and Conservation 28:2295–2317.

Motzkin, G., D. Foster, A. Allen, J. Harrod, and R. Boone. 1996. Controlling site to evaluate history: vegetation patterns of a New England sand plain. Ecological Monographs 66:345–365.

Motzkin, G., P. Wilson, D. R. Foster, and A. Allen. 1999. Vegetation patterns in heterogeneous landscapes: the importance of history and environment. Journal of Vegetation Science 10:903–920.

Nguyen, H. H., J. Uria-Diez, and K. Wiegand. 2016. Spatial distribution and association patterns in a tropical evergreen broad-leaved forest of north-central Vietnam. Journal of Vegetation Science 27:318–327.

Oliver, C. D. 1978. The development of northern red oak in mixed stands in central New England.

Oliver, C. D., and E. P. Stephens. 1977. Reconstruction of a mixed-species forest in central New England. Ecology 58:562–572.

Orwig, D., D. Foster, and A. Ellison. 2015. Harvard Forest CTFS-ForestGEO mapped forest plot since 2014. Harvard Forest Data Archive: HF253. Available online: http://harvardforest.fas.harvard.edu.

Orwig, D. A., A. A. Barker Plotkin, E. A. Davidson, H. Lux, K. E. Savage, and A. M. Ellison. 2013. Foundation species loss affects vegetation structure more than ecosystem function in a northeastern USA forest. PeerJ 1:e41.

Orwig, D. A., P. Boucher, I. Paynter, E. Saenz, Z. Li, and C. Schaaf. 2018. The potential to characterize ecological data with terrestrial laser scanning in Harvard Forest, MA. Interface focus 8.

Pederson, N. 2005. Climatic Sensitivity and Growth of Southern Temperate Tress in the Eastern US: Implicatins for the Carbon Cycle. Columbia University New York, New York.

Perala, D., and D. Alban. 1994. Allometric biomass estimators for aspen-dominated ecosystems in the Upper Great Lakes. Research Paper NC-314. US Department of Agriculture, Forest Service, North Central Research Station, St. Paul, MN.

Plotkin, J. B., M. D. Potts, N. Leslie, N. Manokaran, J. LaFrankie, and P. S. Ashton. 2000. Species-area curves, spatial aggregation, and habitat specialization in tropical forests. Journal of theoretical biology 207:81–99.

R Core Team. 2013. R: A language and environment for statistical computing.

Rackham, O. 1986. The history of the countryside. Dent London.

Rasband, W. 2012. ImageJ: Image processing and analysis in Java. Astrophysics Source Code Library.

Raup, H. M., and R. E. Carlson. 1941. The history of land use in the Harvard Forest. The Harvard Forest Bulletin 20:4–62.

Réjou-Méchain, M., O. Flores, N. Bourland, J. L. Doucet, R. F. Fétéké, A. Pasquier, and O. J. Hardy. 2011. Spatial aggregation of tropical trees at multiple spatial scales. Journal of ecology 99:1373–1381.

Roussopoulos, P. J., and R. M. Loomis. 1979. Weights and dimensional properties of shrubs and small trees of the Great Lakes conifer forest. Research Paper NC-178. St. Paul, MN: US Dept. of Agriculture, Forest Service, North Central Forest Experiment Station 178.

Rowlands, W. 1941. Damage to even-aged stands in Petersham, Massachusetts by the 1938 hurricane as influenced by stand condition. MF thesis. Harvard University, Cambridge, Massachusetts.

Russell, E. W. B. 1997. People and the land through time: linking ecology and history. Yale University Press.

Shearman, T. M., J. M. Varner, S. M. Hood, C. A. Cansler, and J. K. Hiers. 2019. Modelling post-fire tree mortality: Can random forest improve discrimination of imbalanced data? Ecological Modelling 414:108855.

Simmons, C. S. 1941. Prospect Hill 1941 Soil Survey..

Smith, W. B., and G. J. Brand. 1983. Allometric biomass equations for 98 species of herbs, shrubs, and small trees. Research Note NC-299. St. Paul, MN: US Dept. of Agriculture, Forest Service, North Central Forest Experiment Station 299.

Spurr, S. H. 1956. Forest associations in the Harvard Forest. Ecological Monographs 26:245–262.

Strobl, C., A.-L. Boulesteix, A. Zeileis, and T. Hothorn. 2007. Bias in random forest variable importance measures: Illustrations, sources and a solution. BMC bioinformatics 8:25.

Sullivan, F. B., M. J. Ducey, D. A. Orwig, B. Cook, and M. W. Palace. 2017. Comparison of lidar-and allometry-derived canopy height models in an eastern deciduous forest. Forest Ecology and Management 406:83–94.

Telfer, E. 1969. Weight–diameter relationships for 22 woody plant species. Canadian Journal of Botany 47:1851–1855.

Ter-Mikaelian, M. T., and M. D. Korzukhin. 1997. Biomass equations for sixty-five North American tree species. Forest Ecology and Management 97:1–24.

Thompson, J., N. Brokaw, J. K. Zimmerman, R. B. Waide, E. M. Everham, D. J. Lodge, C. M. Taylor, D. García-Montiel, and M. Fluet. 2002. Land use history, environment, and tree composition in a tropical forest. Ecological Applications 12:1344–1363.

Thompson, J. R., D. N. Carpenter, C. V. Cogbill, and D. R. Foster. 2013. Four centuries of change in northeastern United States forests. PLOS ONE 8:e72540.

Turner, B., W. Clark, R. Kates, J. Richards, and J. Mathews. 1990. The earth as transformed by human action: global and regional changes in the biosphere over the past 300 years.

Van Gemerden, B. S., H. Olff, M. P. Parren, and F. Bongers. 2003. The pristine rain forest? Remnants of historical human impacts on current tree species composition and diversity. Journal of Biogeography 30:1381–1390.

Velázquez, E., I. Martínez, S. Getzin, K. A. Moloney, and T. Wiegand. 2016. An evaluation of the state of spatial point pattern analysis in ecology. Ecography 39:1042–1055.

Wang, X., T. Wiegand, Z. Hao, B. Li, J. Ye, and F. Lin. 2010. Species associations in an old-growth temperate forest in north-eastern China. Journal of ecology 98:674–686.

Wang, X., T. Wiegand, A. Wolf, R. Howe, S. J. Davies, and Z. Hao. 2011. Spatial patterns of tree species richness in two temperate forests. Journal of ecology 99:1382–1393.

Wartluft, J. L. 1977. Weights of the Small Appalachian Hardwood Trees and Components. Res. Pap. NE-366. Upper Darby, PA: US Department of Agriculture, Forest Service, Northeastern Forest Experiment Station. 4p. 366.

Westveld, M. 1956. Natural forest vegetation zones of New England. Journal of Forestry 54:332–338.

Wiegand, T., and K. A. Moloney. 2004. Rings, circles, and null-models for point pattern analysis in ecology. Oikos 104:209–229.

Wiegand, T., and K. A. Moloney. 2014. Handbook of spatial point-pattern analysis in ecology. Chapman and Hall/CRC, Baca Raton, FL.

Young, H. E., J. H. Ribe, and K. Wainwright. 1980. Weight tables for tree and shrub species in Maine. Maine. Life Sciences and Agriculture Experiment Station. Miscellaneous report (USA).

Zimmerman, J. K., T. M. Aide, M. Rosario, M. Serrano, and L. Herrera. 1995. Effects of land management and a recent hurricane on forest structure and composition in the Luquillo Experimental Forest, Puerto Rico. Forest Ecology and Management 77:65–76.

